# The glycosphingolipid inhibitor eliglustat inhibits autophagy in osteoclasts to increase bone mass and reduce myeloma bone disease

**DOI:** 10.1101/2021.02.05.429906

**Authors:** Houfu Leng, Hanlin Zhang, Linsen Li, Shuhao Zhang, Yanping Wang, Adel Ersek, Emma Morris, Erdinc Sezgin, Yi-Hsuan Lee, Yunsen Li, Jianqing Mi, Qing Zhong, Claire Edwards, Anna Katharina Simon, Nicole J. Horwood

**Affiliations:** Kennedy Institute of Rheumatology, University of Oxford, Roosevelt Drive, Oxford, UK, OX3 7FY; Key Laboratory of Cell Differentiation and Apoptosis of Chinese Ministry of Education, Department of Pathophysiology, Shanghai Jiao Tong University School of Medicine, Shanghai, P.R. China; Computational Biology Department, Carnegie Mellon University, Pittsburgh, Pennsylvania 15217, USA; Institutes of Biology and Medical Sciences, Soochow University, Suzhou, P.R. China; Norwich Medical School, University of East Anglia, James Watson Road, Norwich, UK, NR4 7UQ; Nuffield Dept. Of Surgical Sciences, Botnar Research Centre, University of Oxford, Old Road, Oxford, UK, OX3 7LD; Science for Life Laboratory, Department of Women’s and Children’s Health, Karolinska Institute, Solna, Sweden; MRC Weatherall Institute of Molecular Medicine, MRC Human Immunology Unit, OX3 9DS, Oxford, UK; Shanghai Institute of Hematology, State Key Laboratory of Medical Genomics, National Research Center for Translational Medicine at Shanghai, Ruijin Hospital Affiliated to Shanghai Jiao Tong University School of Medicine, Shanghai, P.R. China; Nuffield Department of Orthopaedics, Rheumatology, and Musculoskeletal Sciences, Botnar Research Centre, University of Oxford, Old Road, Oxford, UK, OX3 7LD

## Abstract

Multiple myeloma (MM) is a fatal hematological malignancy, where the majority of patients are diagnosed with, or develop, destructive and debilitating osteolytic bone lesions. Current treatments for MM bone disease such as the bisphosphonate zoledronic acid can result in deleterious side effects at high doses. In this study, eliglustat, an FDA approved glycosphingolipid inhibitor, was shown to reduce MM bone disease in preclinical models of MM. Mechanistically, eliglustat alters the lipid composition and plasma membrane fluidity and acts as an autophagy flux inhibitor in bone-resorbing osteoclasts (OC). Autophagic degradation of the signaling molecule TRAF3 is key step in OC differentiation; this was prevented by eliglustat in OC precursors. In addition, eliglustat works depend on TRAF3 *in vivo*. Furthermore, the combination of eliglustat and zoledronic acid was found to have an additive effect to reduce MM bone disease, suggesting the potential for combination therapies that would allow for drug dose reductions. Taken together, this project identifies a novel mechanism in which glycosphingolipid inhibition reduces osteoclastogenesis via autophagy and highlights the translational potential of eliglustat for the treatment of bone loss disorders such as MM.

**One Sentence Summary:** Translational use of eliglustat as an autophagy inhibitor to limit bone lesions in multiple myeloma.

## Introduction

Multiple Myeloma (MM) is the second most common haematological cancer and is caused by abnormal plasma cell expansion in the bone marrow (BM) (*1*). Left untreated, MM induces signs of end-organ damage including hypercalcemia, renal failure, anemia, and bone complications (*2*). MM is preceded by the plasma cell disorder monoclonal gammopathy of undetermined significance (MGUS), which is characterized by an abnormal increase in monoclonal immunoglobulin secretion and mild bone loss, but with no evidence of lytic bone lesions (*3*). Bone lesions are one of the disease-defining features, as the progression of disease from a premalignant state into active MM is characterized by the development of osteolytic bone disease (*4*). More than 85% of MM patients suffer from some form of skeletal complication including osteopenia, pathological fracture, spinal cord compression and vertebral collapse (*5*). Novel ways to alleviate both the pain and disability caused by these bone lesions without causing severe side effects represent an unmet clinical need.

During MM, the BM is altered to establish a unique supportive microenvironment for tumour growth and the associated bone disease due in part to interactions between MM cells and OCs. This creates the so-called “vicious cycle” (*6*), that further contributes to MM progression and bone disease. MM cells are capable of secreting factors like receptor activator of nuclear factor kappa-B ligand (RANKL) to enhance OC formation (*7*). Engagement of RANKL, the OC differentiation factor, with its cognate receptor RANK leads to the activation of the NF*κ*B signalling pathway and is required for OC formation. The NF*κ*B pathway is regulated by tumour necrosis factor receptor-associated factors (TRAFs). In particular TRAF6 mediates RANKL-induced osteoclastogenesis (*8*) while TRAF3 inhibits it (*9*).

Autophagy is a conserved pathway that maintains eukaryotic cell homeostasis by degrading and recycling cellular waste including protein aggregates and damaged organelles(*10*). During OC differentiation Beclin-1 is upregulated (*11*) and Atg7 knockdown inhibits the expression of key OC proteins such as tartrate-resistant acid phosphatase (TRAP) and cathepsin K (*12*). Additionally, autophagy-related proteins including ATG5, ATG7, ATG4B, and LC3 regulate a number of OC functions including ruffled border formation, secretory function, and bone resorption ability (*13*). In line with this, autophagy inhibitors such as chloroquine (CQ) have been shown to inhibit OC formation and restore bone mass (*9*). Molecularly, TRAF3, as a suppressor of OC differentiation, is degraded by autophagy upon RANKL induction in BM OC precursors (*9*), thus highlighting an important role for autophagy in the OC formation process. Based on these findings, autophagy inhibitors may be applied to treat bone loss diseases in the clinic.

The first-line anti-resorptive reagents to treat MM bone disease are bisphosphonates (BPs). MM research trials have shown that Zoledronic acid (ZA), a nitrogen-containing BP, is superior in decreasing bone lesions compared to non-nitrogen-containing BPs (*14, 15*). However, high doses of ZA as used in MM patients may result in deleterious side effects such as osteonecrosis of the jaw (*16*) and can lead to suppression of bone turnover and increased fracture risk (*17*) in ~4% of patients (*14*). As a consequence, continuation of ZA treatment after 2 years requires an interval or break that jeopardizes clinical outcomes. Therefore, treatment strategies for MM bone disease have been extensively investigated in the past few decades.

Glycosphingolipids (GSLs) are constituents of the plasma membrane that are expressed in varying ratios and combinations in all cells. We have previously shown that GM3, a GSL abnormally expressed in human MM cells, promotes osteoclastogenesis (*18*). GSL inhibitors, such as eliglustat, prevent the synthesis of GSL by inhibition of glucosylceramide synthase (GCS) which converts ceramide to glucosylceramide (GlcCer); an important and rate limiting biochemical step in GSL biosynthesis process (*19, 20*). Eliglustat, as a small molecule, is used for the treatment of long-term Gaucher disease type 1 in adults (*21*). Interestingly, patients with Gaucher’s disease have a 6–50 times increased risk of developing MM or the pre-MM MGUS condition (*22*). Identifying a novel reagent with translational potential brings benefits both to MM patients and the disease-associated socioeconomic burden. Here we demonstrate that eliglustat reduces the bone disease in several pre-clinical models of MM, identify its mode of action as an autophagy inhibitor that prevents TRAF3 degradation in OC, and demonstrate that eliglustat can be used in combination with lower doses of ZA for improved clinical outcomes in MM bone disease.

## Results

### Eliglustat increases trabecular bone volume by inhibiting bone-resorbing OCs in healthy mice

To gain insight into the effect of eliglustat treatment on normal bone mass, C57BL/6 mice were fed with normal chow or eliglustat-containing chow for 19 days prior to collecting the tibiae from the mice for micro-CT analysis. Treatment with eliglustat improved trabecular bone as evidenced by micro-CT reconstruction images (Fig. 1A). Quantification demonstrated a significant increase in bone parameters including bone volume (BV/TV), bone surface (BS), trabecular number (Tb.N) and connective density (Conn.D), and a consequent decrease in trabecular separation (Tb.Sp) and in bone surface density (BS/BV). Both BS and BV increased in the eliglustat group, however, as BV was increased much more than BS there was an overall decrease of BS/BV ratio (Fig. 1B). The increase in trabecular bone was not accompanied by a change in either body weight or bone marrow adipose tissue (BMAT) volume, indicating that eliglustat does not substantially affect the lipids in BMAT (Fig. S1).

**Fig. 1.**
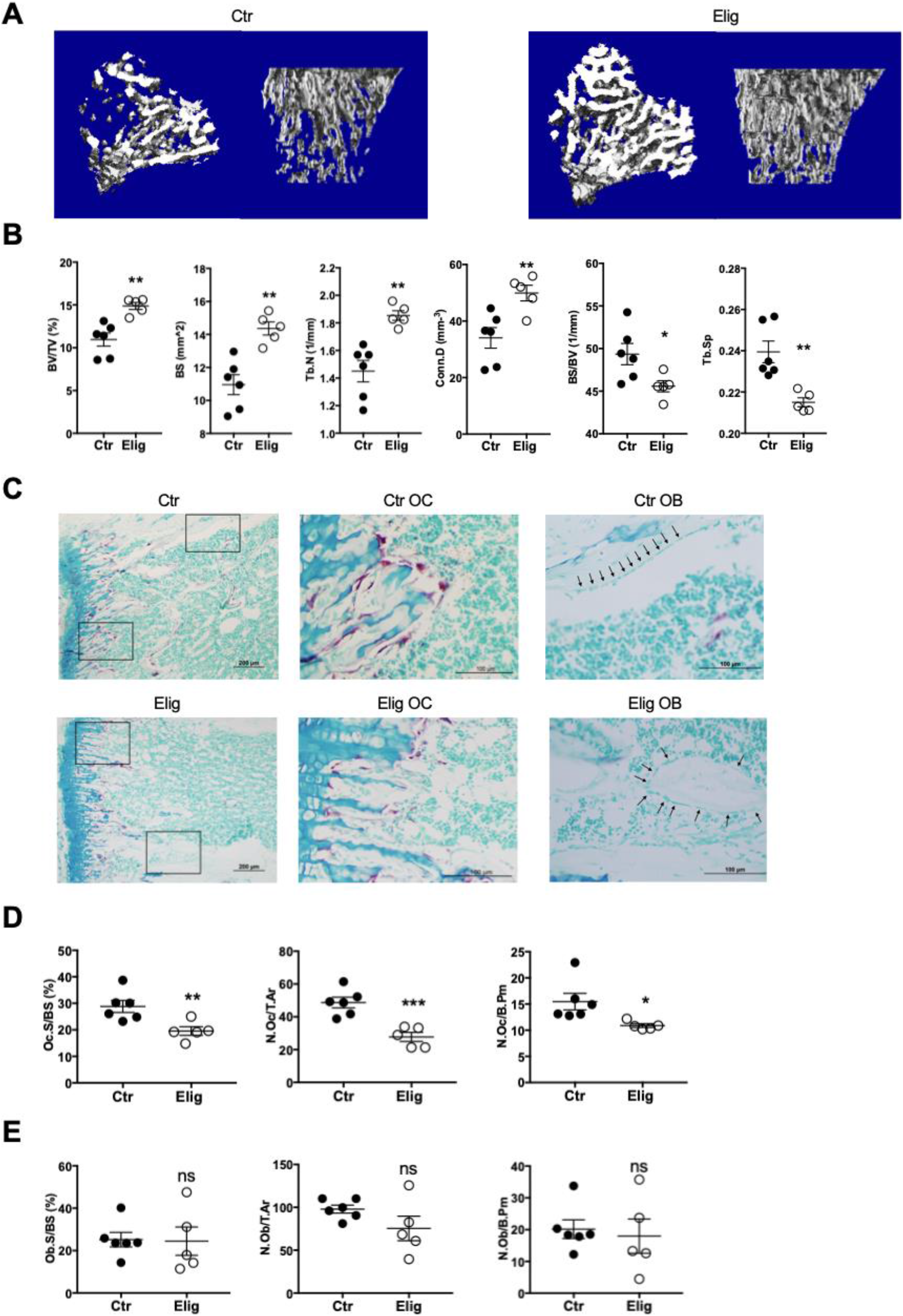
Eliglustat increases trabecular bone in healthy mice by inhibiting OC *in vivo*. (**A**) Representative micro-CT reconstruction images of tibiae from 8-week-old female C57BL/6J mice treated with normal chow (Ctr, *n*=6) or eliglustat (Elig, *n*=5) chow for 19 days. (**B**) Micro-CT analysis of tibiae: BV/TV, BS, Tb.N, Conn.D, BS/BV and Tb.Sp. (**C**) Representative TRAP/0.2% methyl green stained tibial sections showing OCs (red/purple) and OBs (black arrow head) on the endocortical bone surface from each group (left); magnified boxed areas in images on the right. (**D**) Bone histomorphometry quantification of OCs includes the percentage of OC surface over total bone surface (Oc.S/BS), number of OCs over total bone area (N.Oc/T.Ar) and number of OCs per bone perimeter (N.Oc/B.Pm). (**E**) Bone histomorphometry quantification of OB including percentage of OB surface over total bone surface (Ob.S/BS), number of OBs over total bone area (N.Ob/T.Ar) and number of OBs per bone perimeter (N.Ob/B.Pm). *n* = 5-6 as indicated by individual dots, each dot is one mouse. Error bars correspond to SEM. \**P*<0.05, \*\**P*<0.01, \*\*\**P*<0.001. ns means non-significant. Statistical analysis was performed using Student’s *t* test.

To determine whether bone-resorbing OCs or bone-forming osteoblasts (OBs) are the target of eliglustat, their number in trabecular bone was quantified by histomorphometric analysis. Longitudinal cross-sections of tibiae were stained with TRAP to indicate OC. OBs were identified by their typical cuboidal morphology along the bone surface (Fig. 1C). Eliglustat-treated mice tibiae showed significantly lower measurements for OC surface, number and perimeter when compared with naïve control mice (Fig. 1D). However, no difference was observed between untreated control mice and eliglustat-treated mice for OB parameters (Fig. 1E), indicating that the observed increases in trabecular bone were a consequence of OC inhibition rather than an increase in OB.

### Eliglustat decreases MM bone disease in vivo

Eliglustat is primarily used for treating Gaucher’s disease, a rare disease that has an increased risk of MM development (*23, 24*). Gaucher’s patients suffer from osteoporosis/osteopenia and treatment to reduce excess bioactive GSL associated with the disease has been shown to alleviate bone loss(*25*). However, it remains to be determined whether the reduction in bone symptoms is due to the treatment of Gaucher’s disease or if eliglustat acts directly on bone cells, which raised the possibility that eliglustat may be of benefit in the MM bone complications. The 5TGM1 murine model of MM in susceptible C57BL/KaLwRijHsd mice follows a predictable disease course of around 23 days duration as determined by tumor burden in the bone marrow and spleen, the presence of osteolytic bone lesions near the growth plates, and increased serum paraprotein levels (*26*).

To test whether eliglustat could be used as a therapeutic compound under MM conditions, 8-week-old C57BL/KaLwRijHsd mice were injected with saline or 5TGM1-GFP murine MM cells via the tail vein on day 0. Eliglustat-containing chow was given from day 4 until the mice were sacrificed on day 23. Micro-CT reconstruction of tibiae indicated a decrease in trabecular bone in MM-bearing mice compared to saline control group, whilst the eliglustat treated group was significantly protected from MM-induced bone loss (Fig. 2, A and B). Treatment with eliglustat in MM-bearing mice significantly increased BV/TV, BS, Tb.N, Conn.D and Tb.Th, accompanied with a decrease of BS/BV and Tb.Sp (Fig. 2B), confirming that eliglustat inhibits bone catabolic effects. Osteolytic lesions in the cortical bone of 5TGM1-GFP MM-bearing mice were clearly visible by micro-CT reconstruction. Importantly, treatment with eliglustat significantly reduced the number of cortical bone lesions in the tibiae of MM-bearing mice (Fig. 2, C and D).

**Fig. 2.**
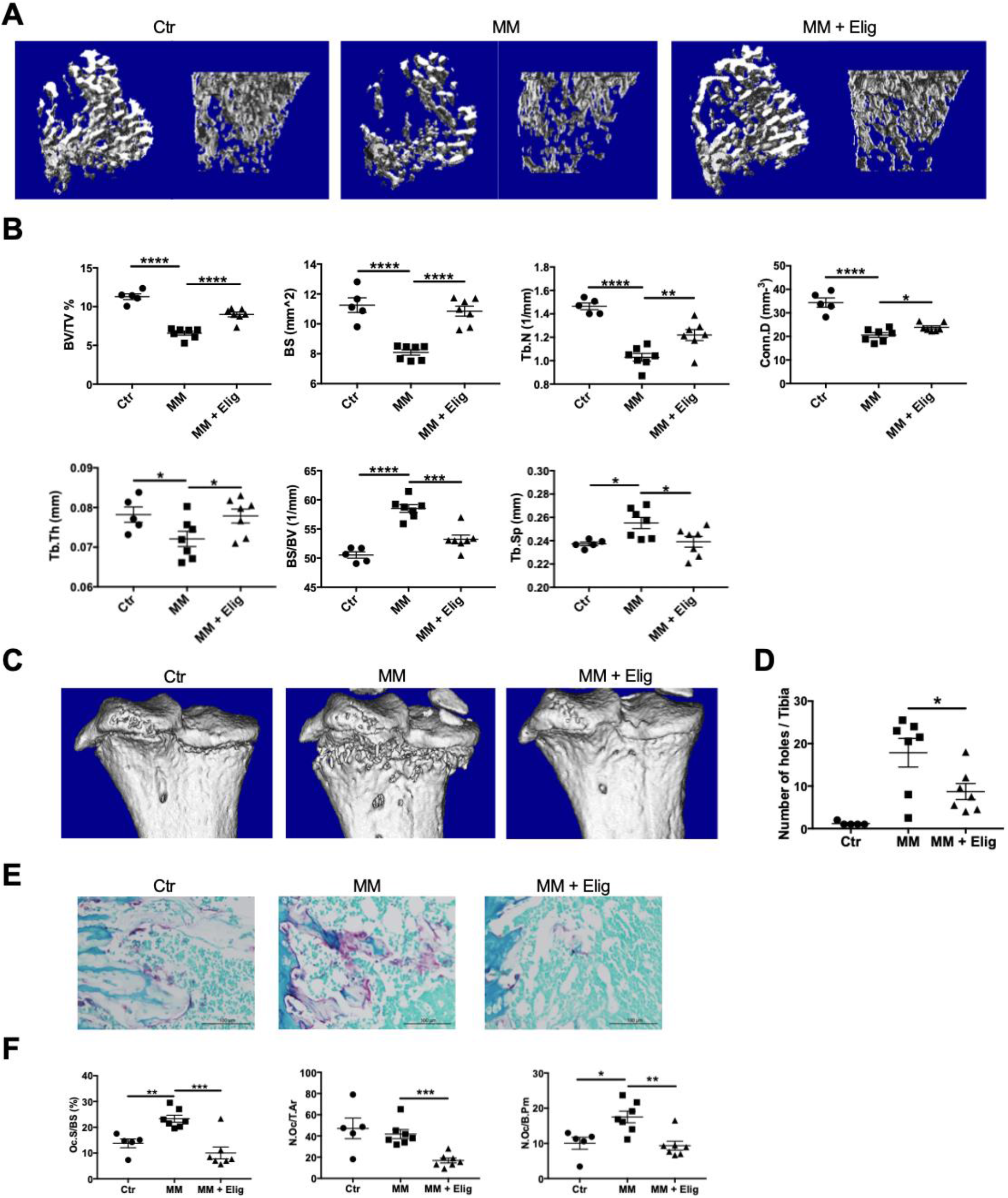
Eliglustat ameliorates 5TGM1-GFP MM cell-induced bone disease. 5TGM1-GFP MM cells were injected to 8-week-old C57BL/KaLwRij mice to generate the MM model. Eliglustat chow was administered from day 4-post tumor injection (until day 23). (**A**) Representative micro-CT reconstruction images of tibiae trabecular bones from naive control (Ctr, *n*=5), 5TGM1-GFP MM bearing mice with normal chow (MM, *n*=7), or MM bearing mice with Eliglustat chow (MM + Elig, *n*=7). (**B**) Trabecular bone parameters in the tibiae were assessed: BV/TV, BS, Tb.N, Conn.D, Tb.Th, BS/BV and Tb.Sp. (**C/D**) Tibiae cortical bone reconstruction (**C**) and the number of cortical bone lesions (**D**). (**E/F**) Representative TRAP/methyl green staining showing red OCs of tibial histological sections, original magnification 40X in (**E)**and the quantification of OCs with indicated parameters in (**F**). Error bars correspond to SEM. \**P*<0.05, \*\**P*<0.01, \*\*\**P*<0.001, \*\*\*\**P*<0.0001. ns, non-significant. Statistical analysis was performed using One-way ANOVA.

As expected, histomorphometric analysis demonstrated a significant increase in TRAP-positive OCs on the endosteal surface in MM-bearing mice compared to control mice while OCs in MM mice exhibited a robust response to eliglustat as shown by decreased parameters including surface, number and perimeter (Fig. 2, E and F). Eliglustat treatment did not change the population of OC precursors as demonstrated by expression of surface markers (CD11b-CD3-B220-CD115+) (*27*) in MM-bearing mice (Fig. S2, A-C) suggesting that eliglustat blocks OC formation rather than inhibiting the generation of OC progenitors.

To determine whether treatment with eliglustat had an effect on MM tumour burden, flow cytometry of ex vivo bone marrow and spleen was undertaken. At the time of sacrifice, there was no difference in GFP+ 5TGM1 MM cell numbers found in BM or spleen between MM-bearing mice and mice treated with eliglustat (Fig. S2, D and F). Systemic tumour burden evaluated by serum paraprotein IgG2b*κ*, a specific marker secreted by MM cells, likewise showed that eliglustat did not reduce tumour burden (Fig. S2G), at least not at this time point. Therefore, eliglustat ameliorates MM-induced bone disease by specifically inhibiting OC without affecting MM tumour burden.

### Eliglustat prevents bone loss in a diet-induced obesity model of MGUS

Patients with Gaucher’s disease have an increased risk of developing the pre-MM condition MGUS (*28, 29*). To expand the potential applications of eliglustat, a well-established high fat diet (HFD) induced murine MGUS model in C57BL/6J was used to evaluate the impact on bone loss (Fig. 3A) (*3*).

**Fig. 3.**
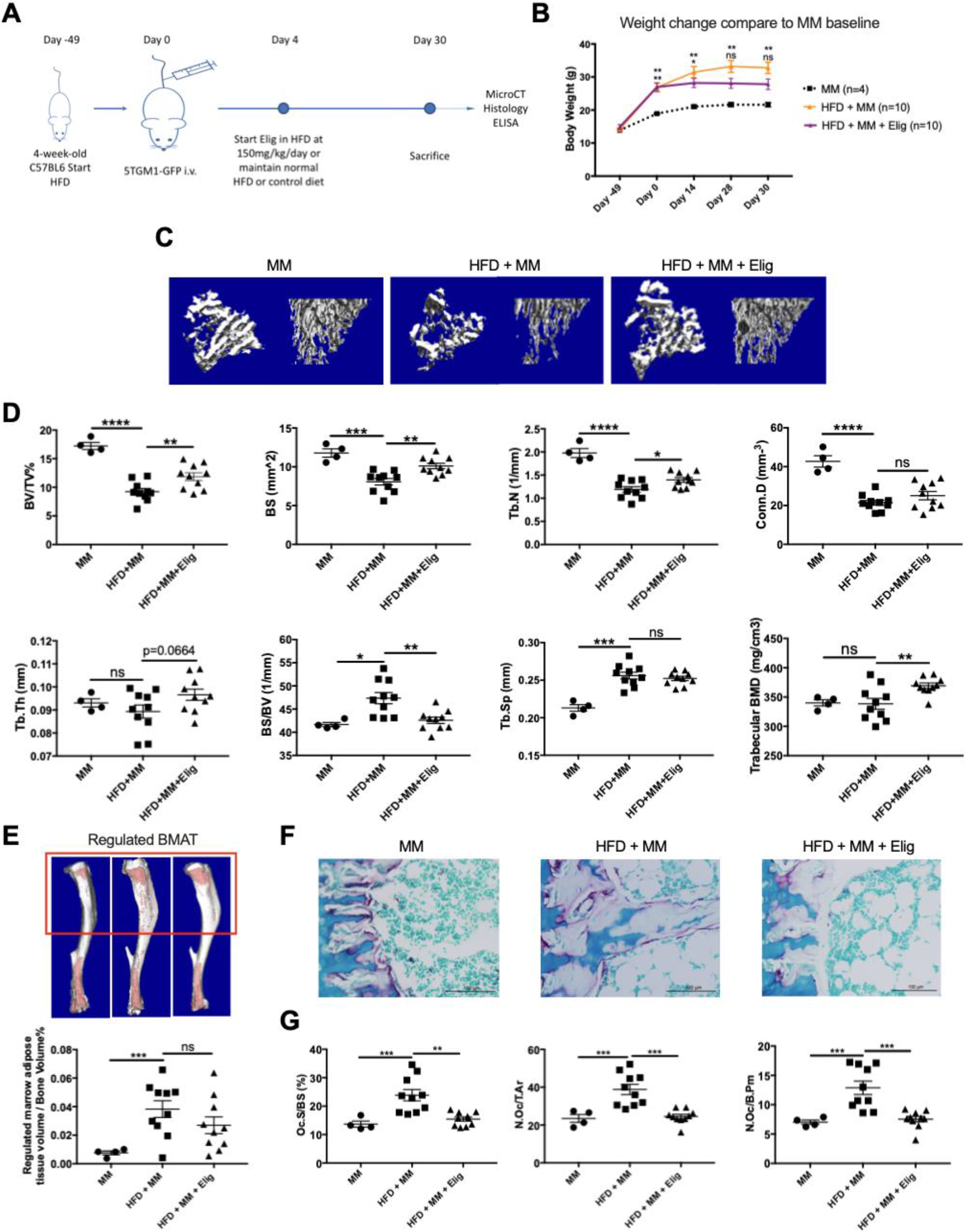
Eliglustat reduces bone loss in a diet-induced obesity MGUS model. (**A**) Schematic overview and experiment design. C57BL6 mice were divided into MM injection with normal diet (MM, *n*=4), MM injection with HFD (MM + HFD, *n*=10) and MM injection with HFD plus eliglustat treatment (MM + HFD + Elig, *n*=10). (**B**) Body weight (g) was measured over the duration of the experiment. (**C**) Representative micro-CT reconstruction images of tibiae trabecular bones from respective groups. (**D**) Micro-CT analysis parameters of tibiae: BV/TV, BS, BS/BV, Tb.Sp, Tb.N, Conn.D, Tb.Th and Trabecular BMD. (**E**) Osmium tetroxide staining to detect rBMAT (in red box) and cBMAT by micro-CT in MM, HFD + MM, and HFD + MM + Elig groups. (**F**) Representative TRAP/methyl green stained tibial histological sections showing red OCs. (**G**) Bone histomorphometry parameters including Oc.S/BS, N.Oc/T.Ar and N.Oc/B.Pm. Error bars correspond to SEM. \**P*<0.05, \*\**P*<0.01, \*\*\**P*<0.001. ns means non-significant. Statistical analysis was performed using One-way ANOVA.

Prior to 5TGM1-GFP MM cell injection, C57BL/6J mice were fed with 42% HFD or control diet for 7 weeks, resulting in a significant increase in body weight in HFD group (Fig. 3, A and B). As previously reported, C57BL/6J mice on a control diet and inoculated with 5TGM1 MM cells showed no signs of tumor growth within the bone marrow or evidence of bone loss (Fig. S3A) (*3*). In addition, 7 weeks’ HFD alone did not change the bone parameters (Fig. S3, B and C). As expected, micro-CT reconstruction images and quantitation showed MM cells injected into HFD mice (MGUS condition) significantly decreased tibial trabecular bone mass compared to the mice placed on normal diet injected with MM cells. Interestingly and in line with the previous figure, eliglustat reversed the bone loss with a significant increase in BV/TV, BS, Tb.N and trabecular BMD without affecting tumor growth (Fig. 3, A, C and D). Moreover, consistent with the data in healthy mice, the BS/BV ratio decreased to control values after eliglustat treatment indicating that bone volume was preserved (Fig. 3D).

Obesity is associated with an increase in BMAT. We have previously demonstrated that marrow adipocytes are elevated in the early stages of MM and are supportive of MM growth and survival (*30*). To determine whether eliglustat impacts marrow adiposity, tibiae were stained with osmium tetroxide and visualized by micro-CT. Quantitative analysis of regulated BMAT (red area within the red rectangle) revealed a significant increase of regulated BMAT in HFD+MM (MGUS) as compared to the MM group (control); eliglustat treatment did not alter regulated BMAT volume indicating that the effects of eliglustat were not due to changes in adiposity in the BM (Fig. 3E). Histological analysis revealed that large adipocytes occupied the BM niche after HFD administration either with or without eliglustat (Fig. 3F). OC parameters by histomorphometric analysis including Oc.S/BS, N.Oc/T.Ar and N.Oc/B.Pm were all significantly increased in HFD+MM (MGUS) group as compared to MM group (control). Treatment with eliglustat significantly decreased OC parameters including OC surface, number and area without affecting OB parameters (Fig. 3G, Fig. S3D).

Taken together, treatment with eliglustat in three in vivo situations; healthy mice, MM susceptible mice and the HFD-induced MGUS murine model, it is clear that osteoclastogenesis was inhibited in all scenarios leading to an investigation of the molecular mechanisms for the inhibition of OC.

### Increased plasma membrane fluidity and altered glycosphingolipidomics following eliglustat treatment

During autophagy, autophagosomes with a double membrane form around portions of the cytoplasm, and then fuse with lysosomes, where their contents are degraded and recycled (*10*). Autophagy is a critical part of osteoclastogenesis and given the important role of GSL in membrane curvature and fusion during autophagy, it was hypothesized that preventing ceramide to glycosphingolipid conversion with eliglustat may block autophagy via an alteration of the lipid environment and membrane organization. To investigate whether eliglustat affects overall lipid organization of OCs, murine RAW264.7 cells were treated with eliglustat for 12 hours, then membrane was stained with NR12S dye and the images were evaluated by spectral imaging (Fig. 4A). The generalized polarization (GP) value reveals how packed the lipids are in the cellular membranes (*31*). We observed that eliglustat significantly decreased the GP value (Fig. 4B), indicating a higher fluidity of the plasma membrane.

**Fig. 4.**
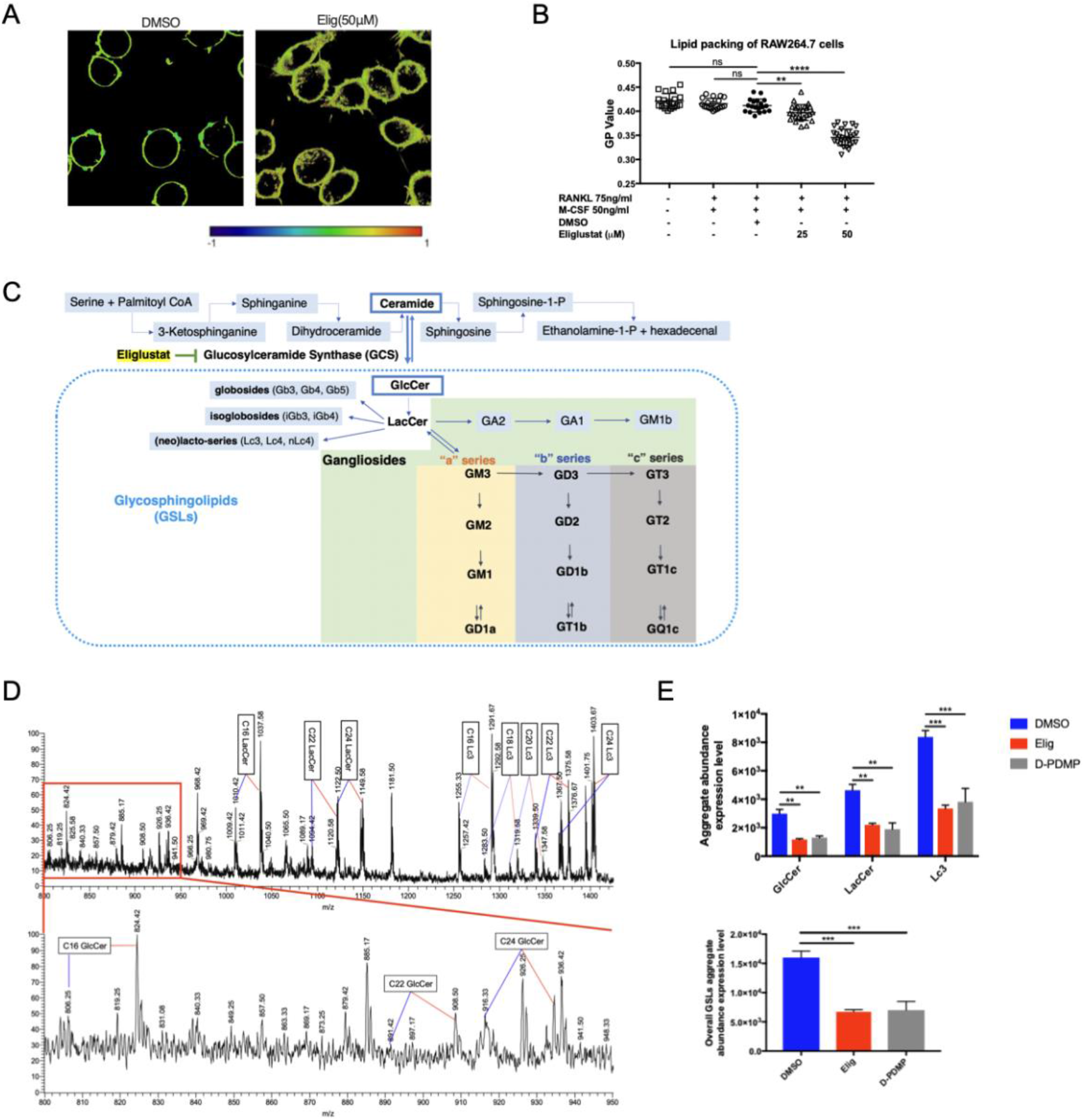
Eliglustat alters the lipid composition in RAW264.7 cells. (**A**) Membrane fluidity during early OC formation process. RAW264.7 cells were treated with 75 ng/ml RANKL and 50 ng/ml M-CSF together with Eliglustat or DMSO for 12 hours. Eliglustat was added to groups at the indicated dose of 25 μM or 50 μM. Representative images of the cells from DMSO group and Eliglustat group at 50μM. Histograms of GP distribution illustrate ordered red end (1) and disordered blue end (−1). (**B**) The quantification for GP value was evaluated. (**C**) Schematic illustrating the pathway of lipid and GSL metabolism. Eliglustat is believed to block the GlcCer formation from Ceramide, which subsequently blocks the formation of LacCer, Gangliosides and other GSLs (including globosides, isoglobosides, and [neo]lacto-series). (**D**) LTQ-ESI-MS glycosphingolipidomics indicating the profile of RAW264.7 cell from DMSO group and eliglustat group (50μM, 12 hours) with 1:1 sample mix to indicate the change of specific GSL composition (green arrow indicates DMSO, red arrow indicates eliglustat) (*m/z* means *mass charge ratio*). (**E**) GlcCer, LacCer and Lc3 aggregate abundance in control, Eliglustat of D-PDMP groups. Error bars correspond to SEM. \*\**P*<0.01, \*\*\*\**P*<0.0001, ns means non-significant. Statistical analysis was performed using One-way ANOVA.

Next, to investigate the lipid composition profile of RAW264.7 cells after eliglustat treatment we used LTQ-ESI-MS mass spectra glycosphingolipidomics to identify various GSLs (Fig. 4C). Following treatment by eliglustat at 50 *μ*M for 12 hours, RAW264.7 cells showed a reduction of GlcCer (cluster of molecular ion peaks between *m/z* 806 and 934), LacCer (cluster of molecular ion peaks between *m/z* 1010 and 1149) and Lc3 from (neo)lacto-series (cluster of molecular ion peaks between *m/z* 1255 and 1403) (Fig. 4, D and E). D-PDMP, a known inhibitor of GCS (*32*), similarly reduced aggregate abundance expression on RAW264.7 cells and served as positive control (Fig. 4E). In summary, eliglustat significantly inhibited GlcCer, LacCer and Lc3 which are all downstream of GCS in RAW264.7 cells, which is likely to be the cause of increased fluidity.

### Eliglustat is a novel autophagy inhibitor that prevents autolysosomal degradation

It has been reported that the degradation of TRAF3 in RANKL-induced OC formation is mediated by autophagy machinery (*9*). Therefore, we investigated whether eliglustat works as an autophagic inhibitor to cause TRAF3 accumulation in OCs. Upon eliglustat treatment, increased accumulation of the autophagosomal and autolysosomal marker LC3-II was observed in RAW264.7 cells and primary BM cells (Fig. 5A); this pattern was similar to treatment using the autophagic flux inhibitor BafilomycinA1 (BafA1; Fig. 5A). BafA1 does not increase LC3-II levels in eliglustat treated RAW264.7 cells any further which indicates that both drugs have an effect late in the autophagic pathway, i.e the lysosomal degradation step. However, this was not observed in primary BM cells and could be due to distinct sensitivity of these two cell types to eliglustat. The inhibition of autophagic flux of RAW264.7 cells has already reached a plateau when eliglustat is administered at 50 μM. Therefore, BafA1 cannot further increase the LC3II level. In contrast, 50 μM eliglustat may not fully block autophagic flux in primary BM cells and thus a further increase upon BafA1 treatment can still be observed. Importantly, p62 protein expression was increased in both RAW264.7 cell and primary cells, which indicates eliglustat is likely to be an autophagy inhibitor.

**Fig. 5.**
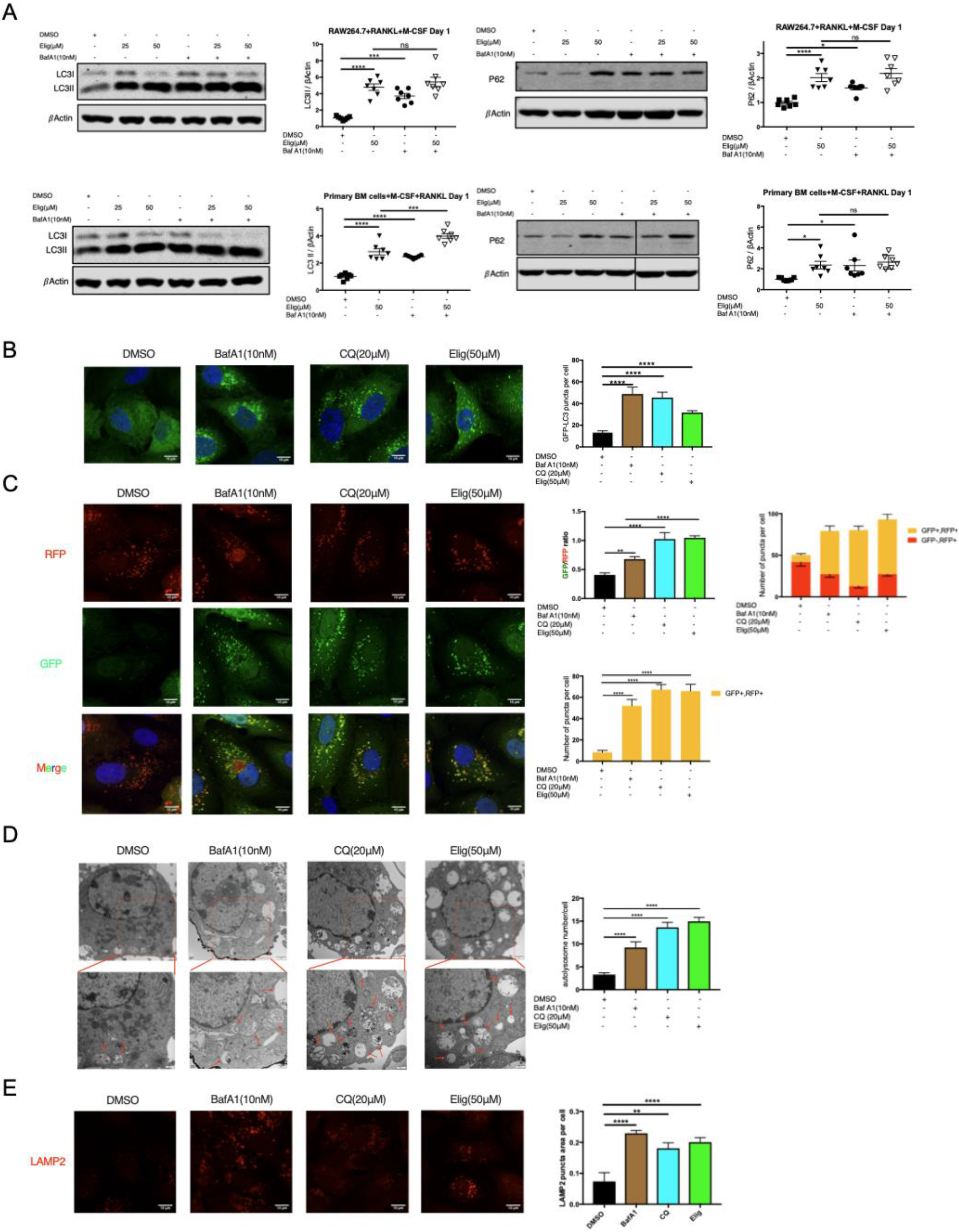
Eliglustat is an autophagy inhibitor. (**A**) RAW264.7 cells and BM cells were treated with RANKL and M-CSF either with or without Eliglustat for 24 hours. 10 nM BafA1 was added 2 hours before protein harvest. Representative western blot showing LC3II protein level and p62 protein level after eliglustat or BafA1 treatment during OC formation process. LC3II and p62 level were quantified. Lanes were run on the same gel, but were non-contiguous. (**B**) BafA1, CQ and eliglustat increased LC3 puncta in U2OS cell transfected with GFP-LC3 plasmid after 2 hours treatment. (**C**) Eliglustat increased GFP/RFP ratio and GFP+RFP+ puncta number in RFP-GFP-LC3 transfected U2OS cell after 6 hours treatment. (**D**) Electron microscope images indicated eliglustat induced autolysosome (red arrow) formation in RAW264.7 cells after 2 hours treatment (25,000X). (**E**) Quantification of the LAMP2 puncta area per U2OS cell (arbitrary units). CQ and BafA1 served as positive control in above experiments. Error bars correspond to SEM. \**P*<0.05, \*\**P*<0.01, \*\*\**P*<0.001, \*\*\*\**P*<0.0001, ns means non-significant. Statistical analysis was performed using One-way ANOVA.

p62 is an autophagic receptor that attracts the cargo into the autophagosomal lumen. Like LC3 it also gets degraded when the autophagsomal content is delivered to the lysosome. We found that p62 degradation was inhibited by BafA1 as expected and by eliglustat in both the RAW264.7 cell line and in primary OC derived from BM (Fig. 5A). Consistently, eliglustat caused significant accumulation of GFP-LC3 puncta in U2OS cells (Fig. 5B).

Both increased autophagosome formation and blocked autophagic flux at the lysosome can lead to accumulation of LC3-II and LC3 puncta. To further distinguish these two possibilities, the tandem-tagged RFP-GFP-LC3 reporter system was used to differentiate whether the accumulated LC3 puncta are autophagosomes or autolysosomes. In this system, GFP is quenched upon autophagic delivery to the acidic lysosome, whereas RFP is more stable in lysosomes. Thus, functional autolysosomes accumulate RFP+ GFP- LC3 (red puncta), and autophagosomes or non-acidic (dysfunctional) autolysosomes accumulate RFP+GFP+ LC3 (yellow puncta) (*33*). As described previously, both BafA1 and CQ that block the fusion of the autophagsomes with lysosomes show an increase in autophagosomes or dysfunctional autolysosomes. Similarly, eliglustat treatment led to increased yellow puncta and reduced red puncta (Fig. 5C), indicating that the accumulation of autophagosomes (or dysfunctional autolysosomes) is associated with reduced lysosomal degradation. Thus, eliglustat blocked autophagic flux via inhibiting lysosomal degradation. This is consistent with the finding that the autophagy substrate p62 and LC3-II itself were accumulated upon eliglustat treatment (Fig. 5A).

Furthermore, transmission electronic microscopy (TEM) revealed that eliglustat caused the accumulation of single-membrane autolysosomes rather than double-membrane autophagosomes in RAW264.7 cells (Fig. 5D), similar to the effect of CQ and BafA1 treatment. Eliglustat also significantly increased the area of LAMP2 puncta, a lysosomal marker, suggesting an increase of lysosomes and/or autolysosomes (Fig. 5E). Taken together, these data support a role for eliglustat as an autophagy inhibitor that blocks autophagic flux by inhibiting autolysosomal degradation.

### OC inhibition with Eliglustat depends on TRAF3

To investigate how the inhibition of autophagy impacts OC differentiation, it was first confirmed that eliglustat inhibits OC differentiation *in vitro* using RAW264.7 cells differentiated with M-CSF and RANKL. Using a dose range below or equivalent to that used clinically (50*μ*M) (*24*), eliglustat inhibited TRAP-positive OC formation in a dose-dependent manner after 5 days of culture (Fig. 6A), as quantified by the number and the area of OCs, as well as the number of nuclei per OC (Fig. 6B). In contrast, cell viability was not affected (Fig. 6B). Consistently, eliglustat also inhibited M-CSF and RANKL-dependent OC formation using OC precursors derived from primary murine BM cells (Fig. S4A).

**Fig. 6.**
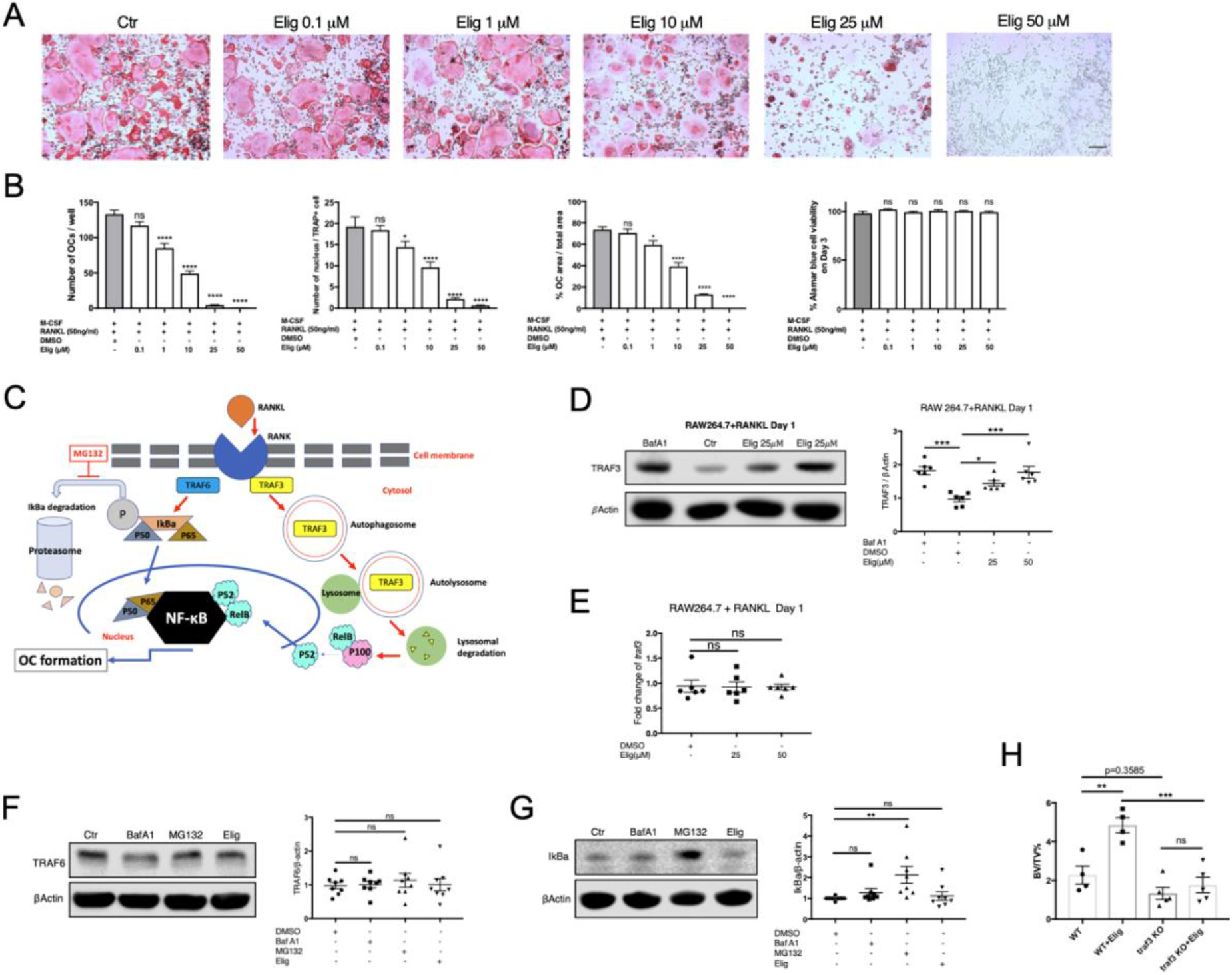
Eliglustat inhibits OC formation in a TRAF3-dependent manner. (**A**) RAW264.7 cells were differentiated into OCs with 50 ng/ml M-CSF and 75 ng/ml RANKL. Different doses of Eliglustat (0.1, 1, 10, 25, 50 μM) were present throughout the culture period and OCs were identified by TRAP staining on day 5. Scale bar represents 200μm. (**B**) Number of OCs per well, number of nucleus per TRAP positive cell, % OC area over total area and Alamar blue viability assay were quantified. (*n*=3) (**C**) Schematic graph illustrating that the binding of RANKL to its receptor activates the TRAF6-dependent canonical NF*κ*B pathway, which leads to proteasome-mediated I*κ*Bα degradation and then nuclear translocation of p65 and p50 whilst in the non-canonical NF *κ* B pathway, binding downregulates TRAF3 via autophagy/lysosome-mediated degradation, which induces nuclear translocation of p52 and RelB. (**D/E**) RAW264.7 cells were treated with M-CSF, RANKL, and eliglustat for 1 day. TRAF3 protein levels were quantified by Western blot and mRNA level was quantified by qRT-PCR. Treatment with BafA1 was for the last 2 hours. (**F/G**) Primary murine BM cells were treated with RANKL together with BafA1 or MG132 or eliglustat for 2 hours. TRAF6 (**F**) and I*κ*Bα (**G**) levels were quantified by Western blot. (**H**) Lethally irradiated recipient mice were reconstituted with littermate (WT) or myeloid specific TRAF3 knockout (LysM-Cre+, Traf3 ^fl/fl)^ bone marrow cells. After 19 days of treatment with eliglustat, bone volume of tibiae was quantified by micro-CT. Data represented as mean ± SEM. \**P*<0.05, \*\**P*<0.01, \*\*\**P*<0.001, \*\*\*\**P*<0.0001, ns, non-significant. Statistical analysis was performed using One-way ANOVA.

Binding of RANKL to the RANK receptor triggers OC formation via the canonical and non-canonical NF-*κ*B pathways (Fig. 6C). In the canonical NF-*κ*B pathway, the activation of the adaptor protein TRAF6 leads to proteasome-mediated I*κ*Bα degradation (*34*) followed by nuclear translocation of p65 and p50; in the non-canonical NF-*κ*B pathway, the adaptor protein TRAF3 is degraded in an autophagosome/lysosome-dependent mechanism (*9*), that induces nuclear translocation of p52 and RelB (*35*). Both pathways are essential for OC formation. To test whether either, or both, pathways were affected by eliglustat, the protein levels of TRAF6 and TRAF3 were measured. Both eliglustat and BafA1 led to the accumulation of TRAF3 protein in RANKL-stimulated RAW264.7 cells (Fig. 6D), whereas traf3 mRNA was not affected (Fig. 6E). This indicates that eliglustat may block TRAF3 lysosomal degradation elicited by RANK activation. Eliglustat-induced TRAF3 accumulation was also observed in primary murine BM cell-derived OCs (Fig. S4B). As expected RANKL up-regulated TRAF6 in murine primary BM cells (*36*) (Fig. S4C), however, this was not altered by eliglustat treatment (Fig. 6F). As previously published, BafA1 (*37*),(*38*) and the proteasomal inhibitor MG132 (*39, 40*), did not affect TRAF6 levels. Additionally, MG132 prevented I *κ* Bα degradation as previously described (*9*), however, eliglustat did not alter I*κ*Bα protein levels (Fig. 6G) indicating that eliglustat does not regulate the canonical NF-*κ*B pathway.

To confirm the role of TRAF3 in OC responses to eliglustat *in vivo*, BM chimeric mice were generated using BM donor cells from LysM-cre; TRAF3^fl/fl^ mice that harbor a myeloid specific deletion of TRAF3 (*9*) (Fig. S4D and E). Chimeric mice were fed eliglustat-containing or control diet for 19 days prior to collection of the tibiae for micro-CT analysis. As previously demonstrated (Fig. 1), eliglustat administration led to a two-fold increase of BV/TV% in mice that received wild-type BM donor cells, however this was no longer apparent in chimeric mice that received BM from TRAF3 deleted mice (Fig. 6H and S4F), indicating that eliglustat suppresses OC formation in a TRAF3-dependent manner.

### Eliglustat combined with ZA has a superior effect compared to ZA alone

ZA is a widely used bisphosphate for the prevention of bone loss in MM bone disease and osteoporosis. By inhibiting the enzyme farnesyl diphosphate synthase, ZA leads to OC apoptosis via blockade of the mevalonate pathway (*41–44*), whilst our data show that eliglustat inhibits OC during the differentiation process. Therefore the efficacy of a combination strategy using these two OC inhibitors was evaluated to demonstrate if there was any potential additive effect.

C57BL/KaLwRijHsd mice were divided into 5 groups: untreated control mice (Ctr); 5TGM1-GFP MM-bearing mice (MM); 5TGM1-GFP MM-bearing mice treated with ZA (MM+ZA; 0.01mg/kg once); MM-bearing mice treated with eliglustat (MM+Elig); or MM-bearing mice treated with both ZA and eliglustat (MM+ZA+Elig) (Fig. 7A). Eliglustat alone and ZA alone both reduced the 5TGM1-GFP cell-induced trabecular bone loss in the tibiae, however the combination strategy was superior to the single use of each compound (Fig. 7B and C). Treatment with the combination of eliglustat and ZA resulted in a 2-fold increase in BV/TV when compared with MM–bearing mice and a 1.5-fold increase when compared with mice treated with eliglustat alone or ZA alone. At the doses used, the effectiveness of eliglustat was similar to ZA with no statistically significant difference between these two groups for BV/TV (Fig. 7C). Importantly, the combination strategy increased BS, BS/BV and Tb.N parameters when compared with ZA given alone, whilst Tb.Sp was significantly decreased (Fig. 7C). Consistent with the finding from trabecular bone, eliglustat and ZA reduced the number of osteolytic lesions in cortical bone to a similar level, whilst the combination of the two compounds significantly decreased the number of holes compared to single use of eliglustat or ZA (Fig. 7D and E).

**Fig. 7.**
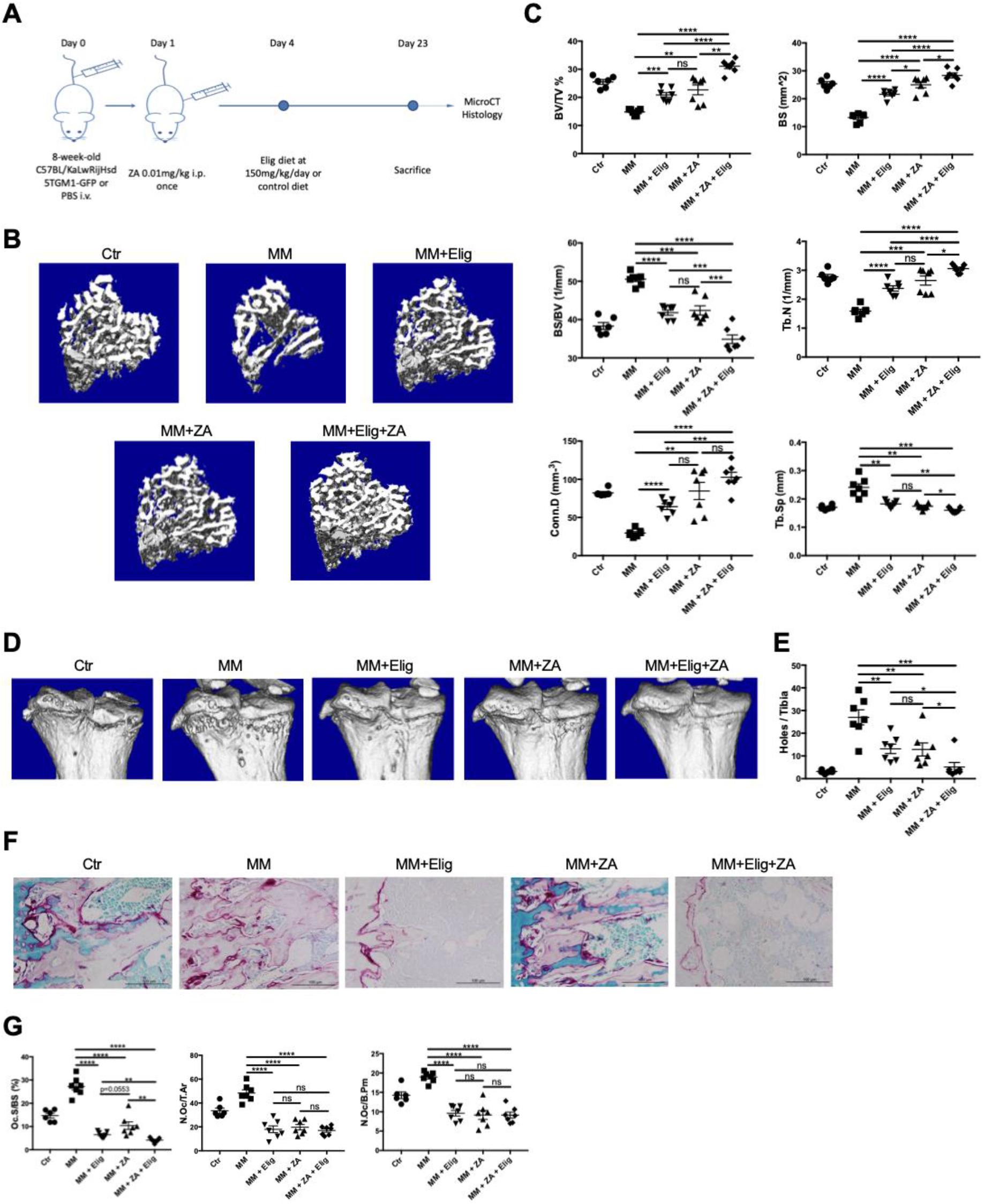
Eliglustat combined with ZA reduces MM bone disease with greater effect than either agent alone. (**A**) Cartoon illustrating the time line and experimental design in investigating how eliglustat and ZA impact on bone measurements in 5TGM1-GFP MM bearing male mice. (**B**) Representative images of 3D micro-CT reconstruction of proximal tibiae from each group, including naive control (Ctr, *n*=6), MM bearing mice (MM, *n*=7), MM bearing mice with eliglustat chow (for 19 days) (MM + Elig, *n*=7), MM bearing mice with single dose ZA injection (0.01 mg/kg) (MM + ZA, *n*=7), or, MM bearing mice with single dose ZA injection (0.01 mg/kg) and eliglustat chow (for 19 days) (MM + ZA + Elig, *n*=7). (**C**) Dot plots of BV/TV, BS, BS/BV, Tb.Sp, Tb.N and Conn.D from micro-CT analysis for each group. (**D**) Representative 3D reconstruction images of proximal cortical bone from each group. (**E**) Bone lesions (holes) on the cortical bones from each mouse were manually counted. (**F**) Representative TRAP/0.2% methyl green stained tibial sections showing red OCs on the endocortical bone surface from each group. (**G**) Bone histomorphometry parameters including Oc.S/BS, N.Oc/T.Ar and N.Oc/B.Pm. Error bars correspond to SEM. \**P*<0.05, \*\**P*<0.01, \*\*\**P*<0.001, \*\*\*\**P*<0.0001. ns means non-significant. Statistical analysis was performed using One-way ANOVA.

Histomorphometric analysis revealed that eliglustat alone, ZA alone and the combination strategy all significantly decreased OC number (N.Oc/T.Ar and N.Oc/B.Pm) compared with the MM-bearing group (Fig. 7F and G). The combination strategy was superior to single use of each compound in inhibiting OC surface (Oc.S/BS) (Fig. 7G). Importantly, all OC parameters in the three treatment groups were significantly lower than non-tumour control (Ctr). Neither eliglustat nor ZA demonstrated a significant effect on OB parameters (Fig. S7A). Thus, the combination of eliglustat and ZA may provide a better clinical strategy than ZA alone for MM patients and other forms of bone loss.

## Discussion

As MM bone disease is primarily caused by the increased numbers and over-activation of bone-resorbing OC, the development of compounds that can inhibit OC formation or function are of great clinical therapeutic importance. Current strategies for OC inhibition involve the use of ZA with limitations to the duration of therapy leading to the need for other ways to inhibit osteoclastogenesis. In this study, eliglustat was assessed in different murine MM models; the C57BL/KaLwRijHsd MM model and C57BL/6J HFD MM model both exhibit bone loss as a consequence of increased tumor burden.

By inhibiting bone resorbing OC formation, eliglustat prevents further bone loss *in vivo* without affecting OC progenitors, OBs, MM cells, HSCs or cells from myeloid linage. While the vicious cycle of MM would suggest that OC inhibition should lead to an indirect reduction in tumor burden or that a reduction in tumor burden is required to decrease OC, the short duration of the model may prevent this from being observed. Indeed, it is not unprecedented to demonstrate an effect on bone disease with no effect tumor burden in murine MM models (*45, 46*). MM-bearing mice with diet-induced obesity have a different lipid metabolism from healthy mice due to excessive adipose tissue. Despite the alterations in lipid metabolism throughout the body, eliglustat successfully improved trabecular bone measurements in MM mice administered with HFD. In addition, ZA, the current first-line bisphosphonate for MM bone disease, had an additive effect in improving bone quality when combined with eliglustat *in vivo*.

Based on the potent effect of eliglustat on OC observed *in vivo*, mechanistic studies were conducted that will underpin future studies in this area. The lipid structure of RAW264.7 cell was more fluid after the administration of eliglustat (Fig. 4A), which can be achieved by reducing cholesterol or increasing the unsaturated lipids in the plasma membrane (*47*). The accumulation of GSLs in cell is known to promote cholesterol accumulation, for instance, GSL inhibitor PDMP enhances cholesterol efflux via the ABCA1/apoA-I pathway(*48*). However, Miglustat, an imino sugar which is a structurally unrelated GSL inhibitor, was unable to stimulate apoA-I-mediated cholesterol efflux(*48*), indicating the complexity of GSL inhibitors. Lipid rafts are a functional membrane units in charge of signal transduction and is composed of cholesterol and saturated lipids (*47*). Therefore, eliglustat may affect lipid rafts and interfere with signaling required for optimal OC formation (*18*). Ha *et al.* discovered that lipid raft domains play an essential role in OC differentiation and affect RANK-TRAF6 signaling (*49*). Likewise, TRAF3 has been located in the lipid raft microdomain by several groups (*50–52*). However, eliglustat did not disrupt canonical NF-*κ*B signaling as TRAF6 was unaffected (Figure 5C and 5F) suggesting that eliglustat may act further downstream to inhibit OC formation. Autophagy is required for the degradation of TRAF3 in the non-canonical NF-*κ*B pathway and hence the effect of eliglustat on the formation and fusion of intracellular vesicles involved in autophagy such as lysosomes or autophagosomes was investigated. Owing to technical difficulties in isolating autophagic or lysosomal membranes in sufficient amount, and the low abundance of GSLs in RAW264.7 cells, the vesicle-specific profile of GSLs by glycosphingolipidomics remains to be thoroughly studied. Interestingly, some GSLs including GD3 (*53*), GM2 and GM3 (*54*) were reported to play an essential role in autophagy. For example, Matarrese *et al.* showed that the ganglioside GD3 contributes to the biogenesis, formation and maturation of autophagic vacuoles (*53*). Therefore, GD3 may be key for autolysosome formation (*53*). As eliglustat inhibits GSL, this may be the mechanism by which eliglustat inhibits autophagy as autophagosome can neither be formed nor matured without GD3.

Due to the accumulation of single-membrane autolysosomes rather than double-membrane autophagosomes in RAW264.7 cells treated with eliglustat (Fig. 5D), we believe that eliglustat is likely to prevent autolysosome degradation thus maintaining TRAF3 levels. Whether this is due to impaired lysosomes remains to be elucidated. For example, lysosomal disruption induced by CQ and other treatments leads to TFEB activation and lysosome biogenesis (*55*). On the other hand, a recent study by Shen *et al.* concluded that another GCS inhibitor (Genz-123346) was an autophagy inducer in HEK293 cells as there was a negative correlation between the increasing dosage of Genz-123346 and a decrease of p62 protein level (*56*). However, our data presented in Fig. 5A did not confirm their findings regarding p62 protein neither in RAW264.7 cell line nor in primary BM cells. Secondly, Genz-123346 caused an accumulation of LC3II protein that did not further increase when treated together with BafA1. This is consistent with our findings in Fig. 5A and suggests that both eliglustat and Genz-123346 are indeed autophagic flux inhibitors as opposed to an inducer. In addition, the findings that eliglustat increased LC3-II levels and mitochondrial mass, and, the co-localization of mitochondria and lysosomes in prostate cancer cells (*32*) are consistent with our data. We observed an increase of LC3-II but also an increase of p62 (Fig. 5A) and when cells were treated with BafA1, there was no further increase of LC3-II indicating that eliglustat blocks autophagic flux. In addition, it was reported that the GSL inhibitor D-PDMP is also an autophagy inhibitor in neuroblastoma cell (*57*).

We noticed that CQ and BafA1, similar to eliglustat, both maintain TRAF3 level and suppress OCs (*9, 58*). However, the FDA approved CQ has considerable ophthalmic side effects and its substitute hydroxychloroquine now used more widely in the clinic had no inhibitory effect on human OC differentiation (*9, 59*). On the other hand, is not an FDA approved drug and its OC inhibition effect was only observed *in vitro*(*58*). Therefore, eliglustat would be a better candidate than either BafA1 or CQ to treat bone lesions in patients.

Our data suggests that inhibition of GSLs and autophagy leading to TRAF3 accumulation is how eliglustat inhibits OC formation and minimizes bone loss. It is nevertheless possible that eliglustat blocks the fusion step of single-nuclei macrophage to multi-nuclei OC as GSLs are present in the plasma membrane. However, cell fusion takes place 3 days after RANKL treatment (*9*), while TRAF3 protein level is already increased in less than 1 day. While fusion resulting in multi nuclei OC may be inhibited by eliglustat, this may happen in addition to the effect on TRAF3 signaling observed here.

We determined that eliglustat could work with ZA to prevent osteoclastogenesis and provide extra bone protection in MM mice by using ZA (0.01 mg/kg) at a ten-fold lower concentration than previous reports (*45*) (Fig. 7) thereby allowing for further increases in bone volume to be observed. The combination of ZA and eliglustat resulted in significantly greater preservation of bone mass than either drug alone. This finding is particularly important as high dose ZA is known to lead to the devastating complication of osteonecrosis of the jaw in MM patients (*16*). By combining it with eliglustat optimal bone protection can be achieved with lower amounts of ZA leading to a potential clinical strategy that is better than current clinical treatments and thus accelerate eliglustat’s translational use for bone protection into the clinic.

There is also tremendous potential for the translational use of eliglustat as a therapeutic in bone loss diseases. Diseases that involve bone loss such as post-menopausal osteoporosis, breast cancer and prostate cancer metastasis to bone, osteomyelitis and arthritis may all exhibit improvement in bone indices as eliglustat was proven to increase bone volume under healthy conditions and to reduce bone loss in the MM disease state. Thus, eliglustat, a clinically approved drug, could be repurposed for patients who suffer from various bone loss diseases and for whom ZA is not suitable, or where they are no longer able to take BP.

## Materials and Methods

### Study design

The objective of the study was to investigate the possibility of eliglustat to increase bone mass and decrease MM bone disease in several pre-clinical models, and decipher the underlying mechanism. Specifically, this study investigated eliglustat inhibits OC formation by preventing TRAF3 degradation, which is due to eliglustat works as an autophagy inhibitor. Using various *in vitro* assays and *in vivo* murine models, the pathway eliglustat involved during OC formation process is revealed, and the specificity and efficacy of eliglustat is evaluated. (i) healthy murine models were used to investigate which cell does eliglustat affect to modulate bone homeostasis, (ii) MM and MGUS pre-clinical models were applied to evaluate the therapeutic potential of eliglustat in various diseases, (iii) myeloid specific TRAF3 knockout (LysM-Cre+, Traf3 ^fl/fl)^ chimeric mice were used to confirm eliglustat works depend on TRAF3, (iv) eliglustat and ZA were administrated to MM mice to compare the treatment efficacy and improved clinical outcomes. Sample size was determined by the authors based on experimental experience. The exact *n* numbers used in each experiment are indicated in the figure legends. For in vivo experiments, animals were assigned randomly to the experimental and control groups. Animal allocation, data acquisition and data analysis *in vivo* or *ex vivo* were performed in a blinded manner.

### Cell lines and reagents

5TGM1-GFP murine MM cells were cultured in RPMI-1640 supplemented with 10% FBS, 1% Penicillin-Streptomycin (P/S), 1% L-Glutamine, 1% Minimum Essential Medium (MEM) Non-Essential Amino Acids and 1% Sodium Pyruvate in 5% CO_2_ at 37°C. RAW264.7 and U2OS cell lines were purchased from ATCC (LGC, UK) and cultured in DMEM with 10% FBS and 1% P/S.

Eliglustat was stored at −20°C until diluted in DMSO immediately before use in culture medium (0.1–50 μM). The *in vitro* OC formation and TRAP experiments were previously described (*18*). Briefly, BM cells or RAW264.7 cells were differentiated into OCs with 20-50 ng/ml M-CSF and 50-75 ng/ml RANKL in MEM Alpha with 10% FBS and 1% P/S, media was changed every 3 days. TRAP assay was conducted on day 5-7 by using the TRAP kit according to manufacturer’s instruction. The reagents details are listed in Supplementary Table 1.

### Mouse models

Animal experiments were undertaken under UK Home Office Project Licenses 30/3218 and 30/3388 in accordance with the UK Animal (Scientific Procedures) Act 1986. Eliglustat was mixed into either a standard rodent diet, or a 42% high fat contained diet at 0.075% w/w (equals 150mg/kg/day), during manufacture by TestDiet. These special diets and respective control diets were provided by Genzyme Corporation (USA). Mice were intravenously injected with 5TGM1-GFP cells or saline as described (*26, 30*). ZA was obtained from Sigma (1724827-150MG) and commenced on day 1 (0.01 mg/kg intraperitoneally once).

Sex and age matched C57BL/6J wild-type mice and CD45.1^+^ B6.SJL mice were purchased from Charles River, UK, and C57BL/KaLwRijHsd mice were purchased from Harlan, The Netherlands. All mice were fed and housed under specific pathogen-free conditions at the Biological Services Unit, Kennedy Institute of Rheumatology, University of Oxford.

### Serum and tissue processing

Mouse blood sampling was collected by tail vein bleeding using capillary blood collection tubes (16.440, Microvette^®^ CB 300 Z). These 10ul samples were stored on ice for up to 2 hours and centrifuged at full speed for 5 minutes to extract serum (supernatant). On the day of cull, serum was collected by intracardiac puncture. Serum was stored at −80°C until further analysis.

Mouse spleens were passed through 70 μm cell strainers (542070, Greiner Bio-One) with PBS, 0.1% BSA and 2mM EDTA to obtain single-cell suspensions. After centrifugation for 3 minutes at 400g in 4°C, red blood cells were lysed with RBC Lysis Buffer (Sigma, R7757-100ML) for 5 minutes at room temperature. RBC-lysed splenocytes were washed once more with PBS, 0.1% BSA, 2mM EDTA and then used for flow cytometry experiment.

Mouse bone marrow cells were collected from the femur. Bones were crushed with a mortar and pestle in PBS-0.1% BSA-2 mM EDTA and filtered through 70 μm cell strainers, then stored on ice ready to be used for flow cytometry without RBC lysis.

### Flow cytometry

Cells were first stained with fixable Zombie Aqua Live/Dead staining and FcR block in PBS for 20 minutes. Samples were then topped up with PBS-0.1% BSA-2mM EDTA as a wash. After centrifugation for 3 minutes at 400g in 4°C, the samples were stained with surface marker antibodies in PBS-0.1% BSA-2mM EDTA at 4°C for another 20 minutes. The cells were washed once with PBS-0.1% BSA-2mM EDTA. Stained cells were analyzed using four-laser LSR Fortessa X-20 (BD Bioscience). Acquired data was analyzed using FlowJo 10.2.

### Antibodies

All the antibodies employed are listed in Supplementary Table 2.

### Chimera Blood Lineages Analysis

The method was previously described(*60*). Briefly, BM chimera was conducted on 11Gy lethally-irradiated female CD45.1^+^ B6.SJL mice using either BM cells from control CD45.2 female mice or CD45.2 BM cells from OC-specific deletion of *traf3* female mice. The reconstitution efficiency of BM chimera was confirmed by evaluating the peripheral blood CD45.2 myeloid population in the CD45.1 mice 6 weeks post-irradiation. 50*μ*l of blood from tail vein bleeding was collected in heparin-coated tubes and stored on ice. The blood was re-suspended in 500*μ*l pre-cold RBC Lysis Buffer for 5 minutes and washed once with 1ml of PBS-0.1% BSA-2mM EDTA. After centrifugation at 300g for 4 minutes, the supernatant was removed and the cells were stained with antibodies for flow cytometry analysis.

### micro-CT analysis

Tibiae were collected and scanned with the Perkin Elmer Quantum FX scanner, Rigaku micro-computed tomography technology, Caliper Life Science according to the manufacturer’s instructions. The region of interest was defined as 100 slices below the growth plate of proximal tibia and was scanned at energy of 90KV; 200uA; field of view (FOV) 10 for 3 minutes. Scanned data was analyzed by using Analyse 12.0 software and the bone 3D reconstruction images were exported.

### Osmium tetroxide staining for BMAT

Osmium tetroxide stain was used to identify BMAT via micro-CT as previously described(*30*). Briefly, tibiae were transferred into osmium tetroxide solution (Honeywell Fulka, 251755-2ml) for 48 hours at room temperature in the fume hood. Then, the tibiae were washed five times with water and embedded in 1% agarose gel. Finally, tibiae are scanned by micro-CT using 90KV; 200μA; FOV10 for 3 minutes and the acquired images used to overlay with previously scanned ones for the bone architecture.

### TRAP staining and histomorphometry analysis of tibiae

As previously described (*18*), tibiae fixed in 4% PFA at 4°C overnight were decalcified in 10% Ethylenediaminetetraacetic acid (EDTA) decalcification solution for 14 days and then paraffin embedded for sectioning with a Leica RM2235 microtome at 5 μm per section to generate histology slides (VWR 631-1553). Sections were processed and stained with TRAP solution and counterstained with 0.2% methyl green solution (Bioenno Lifesciences, 003027). The Olympus BX51 microscope and osteomeasure (Osteometrics) software were used to quantify OC parameters. OB parameters and BMAT volume were quantified based on the morphology of OB and adipocyte.

### Quantitative PCR

Polymerase chain reaction (PCR) was used to detect the change in gene expression in cells. RNA was extracted using the RNeasy Plus Mini Kit (74134, Qiagen). The concentration of RNA was measured using NanoDrop1000 followed by reverse transcribed the RNA to cDNA using High capacity RNA-to-cDNA Kit (Thermo Fisher, 4387406). Taqman probes (Thermo Fisher) and TaqMan Gene Expression Master Mix (Thermo Fisher, 4369016) were used for the qPCR reactions according to the manufacturer’s instructions. The ViiA 7 Real-Time PCR System was used to conduct the quantitative PCR. The quantification of target mRNAs expression was based on *gapdh*. ΔΔCt method was used in all of the experiments.

### IgG2b*κ* ELISA

To evaluate the systemic 5TGM1-GFP MM tumor burden, serum IgG2b*κ* level was measured by enzyme-linked immunosorbent assay (ELISA) according to the manufacturer’s instructions (Bethyl Laboratories, E90-109).

### Western Blot

Cells were lysed using NP-40 lysis buffer containing proteinase inhibitors (Sigma) on ice. The protein concentration was quantified by BCA Assay (23227, Thermo Fisher). Reducing Laemmli Sample Buffer was then added to the lysate and heated at 95℃ for 10 minutes. 15-30 μg proteins were loaded for SDS-PAGE analysis. NuPAGE Novex 4%–12% Bis-Tris gradient gel with MES running buffer (Thermo Fisher) was used. To improve separation of LC3-I and LC3-II, 15% Tris-HCl gel and SDS running buffer was used. Proteins were transferred to a PVDF membrane (IPFL00010, Merck Millipore) and blocked with 5% skimmed milk- PBST. Membranes were incubated with primary antibodies dissolved in 1% milk overnight and secondary antibodies dissolved in 1% milk with 0.01% SDS for 1 hours and then imaged using the Odyssey CLx Imaging System. Data were analysed using Image Studio Lite.

### Immunofluorescence staining

U2OS cells were transfected to overexpress GFP-LC3 plasmid or RFP-GFP-LC3 plasmid as previously described (*33*) and were treated with eliglustat, BafA1 or CQ. Briefly, cells were fixed using 4% paraformaldehyde for 10 minutes at room temperature and then washed three times with ice cold PBS. The samples were incubated for 10 minutes with PBS containing with 0.1% saponin together with 10% FBS for 60 minutes to permeabilize the cells and block unspecific binding of the antibodies. Then, the cells were incubated in diluted primary antibody in 10% FBS in PBS in a humidified chamber for overnight at 4℃. The solution was decanted and cells were washed with PBS three times. Cells were incubated with secondary antibody for 1 hour in room temperature in the dark. Finally, the cells were washed 2 times and stained with DAPI for 5 minutes in the dark. After a further 3 washes, coverslips were added to the slides with a drop of mounting medium (FluorSave^TM^ Reagent, 345789-20ML) prior to examination under laser scanning confocal microscopy (Olympus Fluoview FV3000 or Leica TCS SP8). Image analysis was performed using MATLAB software.

#### Spectral Imaging

Cell were seeded on glass bottom dishes and cultured for 12 hours before imaging. They were incubated with 1 μM NR12S in L15 media for 5 minutes. Spectral imaging was performed by Zeiss LSM 780 microscope equipped with a 32-channel GaAsP detector array. Laser light at 488 nm was selected for NR12S excitation and the λ-detection range was set between 490 and 691 nm. Images were analyzed with the custom GP plugin of FIJI software as described previously(*61*).

### Sample preparation for glycosphingolipidomics

The sample preparation method was previously described (*62*). Briefly, after methylation, qualitative analysis of fucosylated neutral GSLs was performed in the positive ion mode on the LTQ-ESI-MS mass spectrometer (Thermo Fischer Scientific, Waltham, MA, USA) by using a metal needle for direct infusion of samples dissolved in methanol, with a flow rate of 5 ll/min and at ion spray voltage 3.5 kV, capillary voltage 35 V, capillary temperature 350°C, injection time 100 ms, activation time 30 ms, and isolation width m/z 1.5.

### Transmission electron microscopy

RAW264.7 cells (1 × 10^8^) treated with BafA1 or CQ or eliglustat for 2 hours were followed by fixing with 0.1 M sodium cacodylate buffer solution (pH 7.4) containing 2.5% glutaraldehyde for 1 hours at 4°C. The cells were then scraped and centrifuged at 400 g for 5 minutes at 4°C to collect the pellets. Samples were submitted to the Electron Microscopy Core Facility at Shanghai Jiao Tong University for standard transmission electron microscopy (HITACHI 7650) ultrastructural analyses.

### Statistical analyses

Data analysis was conducted using Prism (GraphPad) software. All data shown are represented as mean ± standard error of the mean (SEM). Two-tailed t-test were applied when comparing two experimental groups. For comparisons between two normally distributed data sets with equal variances, paired or unpaired two-tailed Student’s t-test was applied. For the experiments that contain more than three experimental groups, One-way analysis of variance (ANOVA) and multiple comparisons were used with Tukey’s correction. Paired or unpaired one-way ANOVA was used for multiple comparisons of normally distributed datasets with one variable. To evaluate the statistical significance of the hypothesis being tested, p value was applied. * *P* ≤0.05, ** *P*≤0.01, *** *P* ≤0.001, **** *P* ≤0.0001, ns, not significant. Statistical analyses were performed using GraphPad Prism (San Diego, CA).

## Supplementary Materials

Fig. S1. Eliglustat does not affect body weight and BMAT *in vivo*.

Fig. S2. Eliglustat has no effect on OC progenitors or MM tumor burden in the 5TGM1-GFP MM model.

Fig. S3. Eliglustat does not alter tumor burden. HFD does not change the bone parameters. Quantification of the OBs and cortical OCs.

Fig. S4. Eliglustat inhibits OC formation in a TRAF3-dependent manner.

Fig. S5. Eliglustat combined with ZA does not enhance OB number or area.

Fig. S6. Western blot full scans.

Table S1. Reagents used for cell culture.

Table S2. Antibodies used for flow cytometry, Western blot and confocal.

Table S3. TaqMan^®^ probes.

## Acknowledgments

We thank Xiaoxia Liu and members from Qing Zhong’s lab helped on experiments – it is greatly appreciated. Zhenqiang Yao and Akram Ayoub from Brendan Boyce’s lab in University of Rochester Medical Center provided scientific advice and prepared the TRAF3 KO mouse BM cells. 5TGM1-GFP murine MM cells was provided by B.O. Oyajobi, The University of Texas Health Science Center, San Antonio. Jin Wang and Duohui Jing from Ruijin Hospital, Ge Zhang from Hong Kong Baptist University, Mei Tian from Second Hospital of Zhejiang University, Bin Xie from Kennedy Institute of Rheumatology all provided constructive comments.

## Author Contributions

HL - designed, performed and analyzed experiments; wrote the manuscript

HZ - designed and analyzed experiments; edited the manuscript

LL - designed, performed and analyzed experiments

SZ - performed and analyzed experiments

YW - performed and analyzed experiments

AE - performed and analyzed experiments

EM - performed and analyzed experiments

ES - performed and analyzed experiments

Y-HL - performed experiments

YL - contributed reagents and discussed analyses

JM - contributed reagents and discussed analyses

QZ - designed and analyzed experiments; funding

CE - designed and analyzed experiments; funding; edited the manuscript

KS - designed and analyzed experiments; funding; edited the manuscript

NH - designed and analyzed experiments; funding; wrote the manuscript

## Reference

1. R. L. Siegel, K. D. Miller, A. Jemal, Cancer statistics, 2016. CA Cancer J Clin 66, 7–30 (2016).

2. S. V. Rajkumar et al., International Myeloma Working Group updated criteria for the diagnosis of multiple myeloma. Lancet Oncol 15, e538–548 (2014).

3. S. T. Lwin, S. W. Olechnowicz, J. A. Fowler, C. M. Edwards, Diet-induced obesity promotes a myeloma-like condition in vivo. Leukemia 29, 507–510 (2015).

4. P. R. Greipp et al., International staging system for multiple myeloma. J Clin Oncol 23, 3412–3420 (2005).

5. E. Terpos et al., Soluble receptor activator of nuclear factor kappaB ligand-osteoprotegerin ratio predicts survival in multiple myeloma: proposal for a novel prognostic index. Blood 102, 1064–1069 (2003).

6. M. Abe, K. Hiura, S. Ozaki, S. Kido, T. Matsumoto, Vicious cycle between myeloma cell binding to bone marrow stromal cells via VLA-4-VCAM-1 adhesion and macrophage inflammatory protein-1alpha and MIP-1beta production. J Bone Miner Metab 27, 16–23 (2009).

7. D. F. Quail, J. A. Joyce, Microenvironmental regulation of tumor progression and metastasis. Nat Med 19, 1423–1437 (2013).

8. A. P. Armstrong et al., A RANK/TRAF6-dependent signal transduction pathway is essential for osteoclast cytoskeletal organization and resorptive function. J Biol Chem 277, 44347–44356 (2002).

9. Y. Xiu et al., Chloroquine reduces osteoclastogenesis in murine osteoporosis by preventing TRAF3 degradation. J Clin Invest 124, 297–310 (2014).

10. A. J. Clarke, A. K. Simon, Autophagy in the renewal, differentiation and homeostasis of immune cells. Nat Rev Immunol 19, 170–183 (2019).

11. Y. H. Chung et al., Beclin-1 is required for RANKL-induced osteoclast differentiation. J Cell Physiol 229, 1963–1971 (2014).

12. N. Y. Lin et al., Inactivation of autophagy ameliorates glucocorticoid-induced and ovariectomy-induced bone loss. Ann Rheum Dis 75, 1203–1210 (2016).

13. C. J. DeSelm et al., Autophagy proteins regulate the secretory component of osteoclastic bone resorption. Dev Cell 21, 966–974 (2011).

14. G. J. Morgan et al., Long-term follow-up of MRC Myeloma IX trial: Survival outcomes with bisphosphonate and thalidomide treatment. Clin Cancer Res 19, 6030–6038 (2013).

15. N. D. Modi, S. Lentzsch, Bisphosphonates as antimyeloma drugs. Leukemia 26, 589–594 (2012).

16. A. Badros et al., Natural history of osteonecrosis of the jaw in patients with multiple myeloma. J Clin Oncol 26, 5904–5909 (2008).

17. K. A. Kennel, M. T. Drake, Adverse effects of bisphosphonates: implications for osteoporosis management. Mayo Clin Proc 84, 632–637; quiz 638 (2009).

18. A. Ersek et al., Glycosphingolipid synthesis inhibition limits osteoclast activation and myeloma bone disease. J Clin Invest 125, 2279–2292 (2015).

19. S. Ichikawa, Y. Hirabayashi, Glucosylceramide synthase and glycosphingolipid synthesis. Trends Cell Biol 8, 198–202 (1998).

20. H. Fyrst, J. D. Saba, An update on sphingosine-1-phosphate and other sphingolipid mediators. Nat Chem Biol 6, 489–497 (2010).

21. R. M. Poole, Eliglustat: first global approval. Drugs 74, 1829–1836 (2014).

22. B. E. Rosenbloom, P. Becker, N. Weinreb, Multiple myeloma and Gaucher genes. Genet Med 11, 134 (2009).

23. S. Nair et al., Clonal Immunoglobulin against Lysolipids in the Origin of Myeloma. N Engl J Med 374, 555–561 (2016).

24. E. V. Pavlova et al., Inhibition of UDP-glucosylceramide synthase in mice prevents Gaucher disease-associated B-cell malignancy. J Pathol 235, 113–124 (2015).

25. R. S. Kamath et al., Skeletal improvement in patients with Gaucher disease type 1: a phase 2 trial of oral eliglustat. Skeletal Radiol 43, 1353–1360 (2014).

26. S. T. Lwin, C. M. Edwards, R. Silbermann, Preclinical animal models of multiple myeloma. Bonekey Rep 5, 772 (2016).

27. C. Jacquin, D. E. Gran, S. K. Lee, J. A. Lorenzo, H. L. Aguila, Identification of multiple osteoclast precursor populations in murine bone marrow. J Bone Miner Res 21, 67–77 (2006).

28. M. Arends, L. van Dussen, M. Biegstraaten, C. E. Hollak, Malignancies and monoclonal gammopathy in Gaucher disease; a systematic review of the literature. Br J Haematol 161, 832–842 (2013).

29. N. J. Weinreb, P. K. Mistry, B. E. Rosenbloom, M. V. Dhodapkar, MGUS, lymphoplasmacytic malignancies, and Gaucher disease: the significance of the clinical association. Blood 131, 2500–2501 (2018).

30. E. V. Morris et al., Myeloma Cells Down-Regulate Adiponectin in Bone Marrow Adipocytes Via TNF-Alpha. J Bone Miner Res 35, 942–955 (2020).

31. E. Sezgin et al., Polarity-Sensitive Probes for Superresolution Stimulated Emission Depletion Microscopy. Biophys J 113, 1321–1330 (2017).

32. J. Vykoukal et al., Caveolin-1-mediated sphingolipid oncometabolism underlies a metabolic vulnerability of prostate cancer. Nat Commun 11, 4279 (2020).

33. J. Diao et al., ATG14 promotes membrane tethering and fusion of autophagosomes to endolysosomes. Nature 520, 563–566 (2015).

34. E. Ang et al., Proteasome inhibitors impair RANKL-induced NF-kappaB activity in osteoclast-like cells via disruption of p62, TRAF6, CYLD, and IkappaBalpha signaling cascades. J Cell Physiol 220, 450–459 (2009).

35. A. Oeckinghaus, M. S. Hayden, S. Ghosh, Crosstalk in NF-kappaB signaling pathways. Nat Immunol 12, 695–708 (2011).

36. X. Kong et al., Triterpenoid Saponin W3 from Anemone flaccida Suppresses Osteoclast Differentiation through Inhibiting Activation of MAPKs and NF-kappaB Pathways. Int J Biol Sci 11, 1204–1214 (2015).

37. F. Y. Khusbu et al., Resveratrol induces depletion of TRAF6 and suppresses prostate cancer cell proliferation and migration. Int J Biochem Cell Biol 118, 105644 (2020).

38. S. T. Chan, J. Lee, M. Narula, J. J. Ou, Suppression of Host Innate Immune Response by Hepatitis C Virus via Induction of Autophagic Degradation of TRAF6. J Virol 90, 10928–10935 (2016).

39. S. M. Hindi, A. Kumar, TRAF6 regulates satellite stem cell self-renewal and function during regenerative myogenesis. J Clin Invest 126, 151–168 (2016).

40. Y. H. Wu et al., Bortezomib enhances radiosensitivity in oral cancer through inducing autophagy-mediated TRAF6 oncoprotein degradation. J Exp Clin Cancer Res 37, 91 (2018).

41. A. Hameed, J. J. Brady, P. Dowling, M. Clynes, P. O’Gorman, Bone disease in multiple myeloma: pathophysiology and management. Cancer Growth Metastasis 7, 33–42 (2014).

42. M. V. Dhodapkar et al., Anti-myeloma activity of pamidronate in vivo. Br J Haematol 103, 530–532 (1998).

43. J. R. Berenson et al., Long-term pamidronate treatment of advanced multiple myeloma patients reduces skeletal events. Myeloma Aredia Study Group. J Clin Oncol 16, 593–602 (1998).

44. R. G. Russell, Pharmacological diversity among drugs that inhibit bone resorption. Curr Opin Pharmacol 22, 115–130 (2015).

45. M. M. McDonald et al., Inhibiting the osteocyte-specific protein sclerostin increases bone mass and fracture resistance in multiple myeloma. Blood 129, 3452–3464 (2017).

46. D. J. Heath et al., Inhibiting Dickkopf-1 (Dkk1) removes suppression of bone formation and prevents the development of osteolytic bone disease in multiple myeloma. J Bone Miner Res 24, 425–436 (2009).

47. E. Sezgin, I. Levental, S. Mayor, C. Eggeling, The mystery of membrane organization: composition, regulation and roles of lipid rafts. Nat Rev Mol Cell Biol 18, 361–374 (2017).

48. E. N. Glaros et al., Glycosphingolipid accumulation inhibits cholesterol efflux via the ABCA1/apolipoprotein A-I pathway: 1-phenyl-2-decanoylamino-3-morpholino-1-propanol is a novel cholesterol efflux accelerator. J Biol Chem 280, 24515–24523 (2005).

49. H. Ha et al., Membrane rafts play a crucial role in receptor activator of nuclear factor kappaB signaling and osteoclast function. J Biol Chem 278, 18573–18580 (2003).

50. D. G. Meckes, Jr., N. F. Menaker, N. Raab-Traub, Epstein-Barr virus LMP1 modulates lipid raft microdomains and the vimentin cytoskeleton for signal transduction and transformation. J Virol 87, 1301–1311 (2013).

51. M. Higuchi, K. M. Izumi, E. Kieff, Epstein-Barr virus latent-infection membrane proteins are palmitoylated and raft-associated: protein 1 binds to the cytoskeleton through TNF receptor cytoplasmic factors. Proc Natl Acad Sci U S A 98, 4675–4680 (2001).

52. H. Ardila-Osorio et al., TRAF interactions with raft-like buoyant complexes, better than TRAF rates of degradation, differentiate signaling by CD40 and EBV latent membrane protein 1. Int J Cancer 113, 267–275 (2005).

53. P. Matarrese et al., Evidence for the involvement of GD3 ganglioside in autophagosome formation and maturation. Autophagy 10, 750–765 (2014).

54. C. Bedia et al., GM2-GM3 gangliosides ratio is dependent on GRP94 through down-regulation of GM2-AP cofactor in brain metastasis cells. Sci Rep 9, 14241 (2019).

55. A. Ballabio, J. S. Bonifacino, Lysosomes as dynamic regulators of cell and organismal homeostasis. Nat Rev Mol Cell Biol 21, 101–118 (2020).

56. W. Shen et al., Inhibition of glucosylceramide synthase stimulates autophagy flux in neurons. J Neurochem 129, 884–894 (2014).

57. J. Wei et al., Protective role of endogenous gangliosides for lysosomal pathology in a cellular model of synucleinopathies. Am J Pathol 174, 1891–1909 (2009).

58. S. Zhu et al., Bafilomycin A1 Attenuates Osteoclast Acidification and Formation, Accompanied by Increased Levels of SQSTM1/p62 Protein. J Cell Biochem 117, 1464–1470 (2016).

59. C. K. Lee et al., Effects of disease-modifying antirheumatic drugs and antiinflammatory cytokines on human osteoclastogenesis through interaction with receptor activator of nuclear factor kappaB, osteoprotegerin, and receptor activator of nuclear factor kappaB ligand. Arthritis Rheum 50, 3831–3843 (2004).

60. H. Zhang et al., Polyamines Control eIF5A Hypusination, TFEB Translation, and Autophagy to Reverse B Cell Senescence. Mol Cell, (2019).

61. E. Sezgin, D. Waithe, J. Bernardino de la Serna, C. Eggeling, Spectral imaging to measure heterogeneity in membrane lipid packing. Chemphyschem 16, 1387–1394 (2015).

62. Z. Wang et al., High expression of lactotriaosylceramide, a differentiation-associated glycosphingolipid, in the bone marrow of acute myeloid leukemia patients. Glycobiology 22, 930–938 (2012).

