## Supplemental data for "The glycosphingolipid inhibitor eliglustat inhibits autophagy in osteoclasts to increase bone mass and reduce myeloma bone disease"

**Supplemental Figure 1**

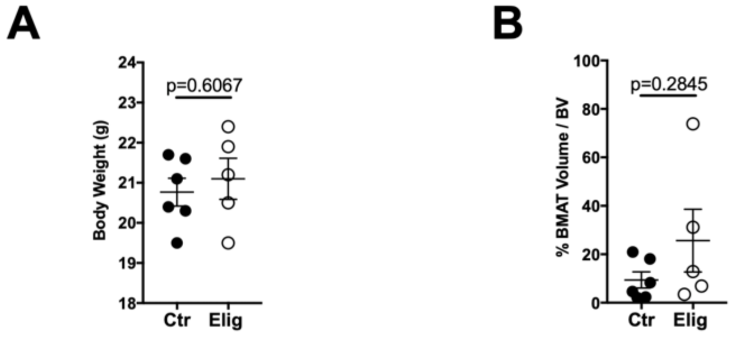

**Fig. S1. Eliglustat does not affect body weight and BMAT *in vivo*.** (**A**) 8-week-old C57BL6 mice were fed on normal chow (Ctr, n=6) or eliglustat chow (150 mg/kg/day, Elig, n=5) for 19 days prior to body weight measurement. (**B**) Bone histomorphometry analysis for the bone marrow adipose tissue (BMAT) volume relative to total BV. Error bars correspond to SEM. Statistical analyses were performed using Student's *t* test.

**Supplemental Figure 2**

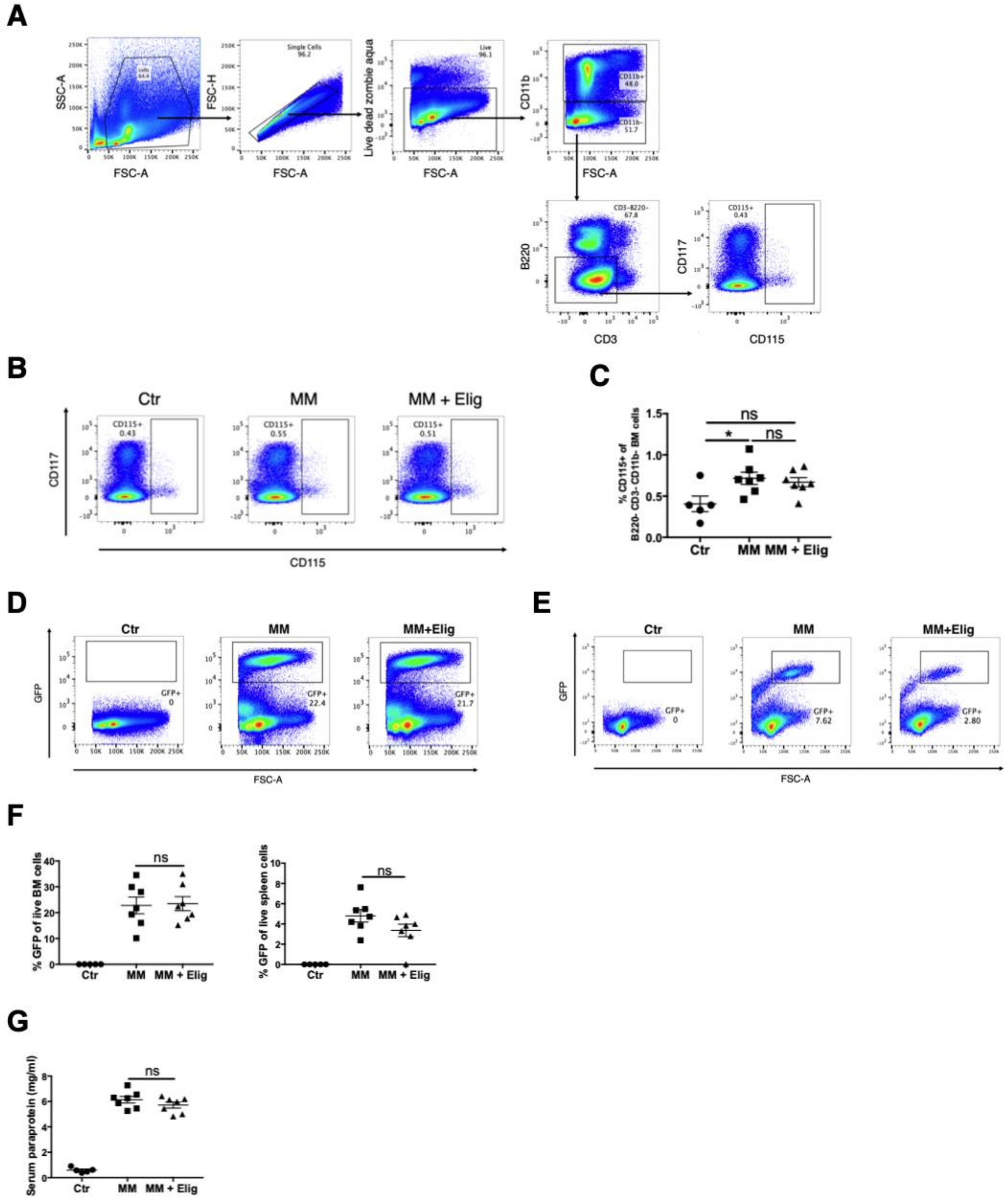

**Fig. S2. Eliglustat has no effect on OC progenitors or MM tumour burden in the 5TGM1-GFP MM model.** BM from female C57BL/KaLwRij mice, either untreated controls (Ctr), myeloma bearing mice with normal chow (MM), or myeloma bearing mice with Eliglustat chow from day 4 post tumour injection (150 mg/kg/d for 19 days, MM + Elig) were collected for flow cytometry analysis. (**A**) Flow cytometry gating strategy of the OC progenitors (CD11b- B220- CD3- CD115+) in BM. (**B-C**) Representative plots of OC progenitors (CD11b- B220- CD3- CD115+) from the indicated groups (**B**) and the quantified percentage (**C**). (**D-F**) Tumour burden in BM (**D**) and spleen (**E**) was assessed by quantifying $GFP^+$ cells using flow cytometry (**F**). (**G**) Serum paraprotein IgG2b*κ* secreted by 5TGM1-GFP cells was quantified using ELISA. Error bars correspond to SEM. *P<0.05, **P<0.01, ***P<0.001, ****P<0.0001. ns, non-significant. Statistical analysis was performed using Student's t-test.

**Supplemental Figure 3**

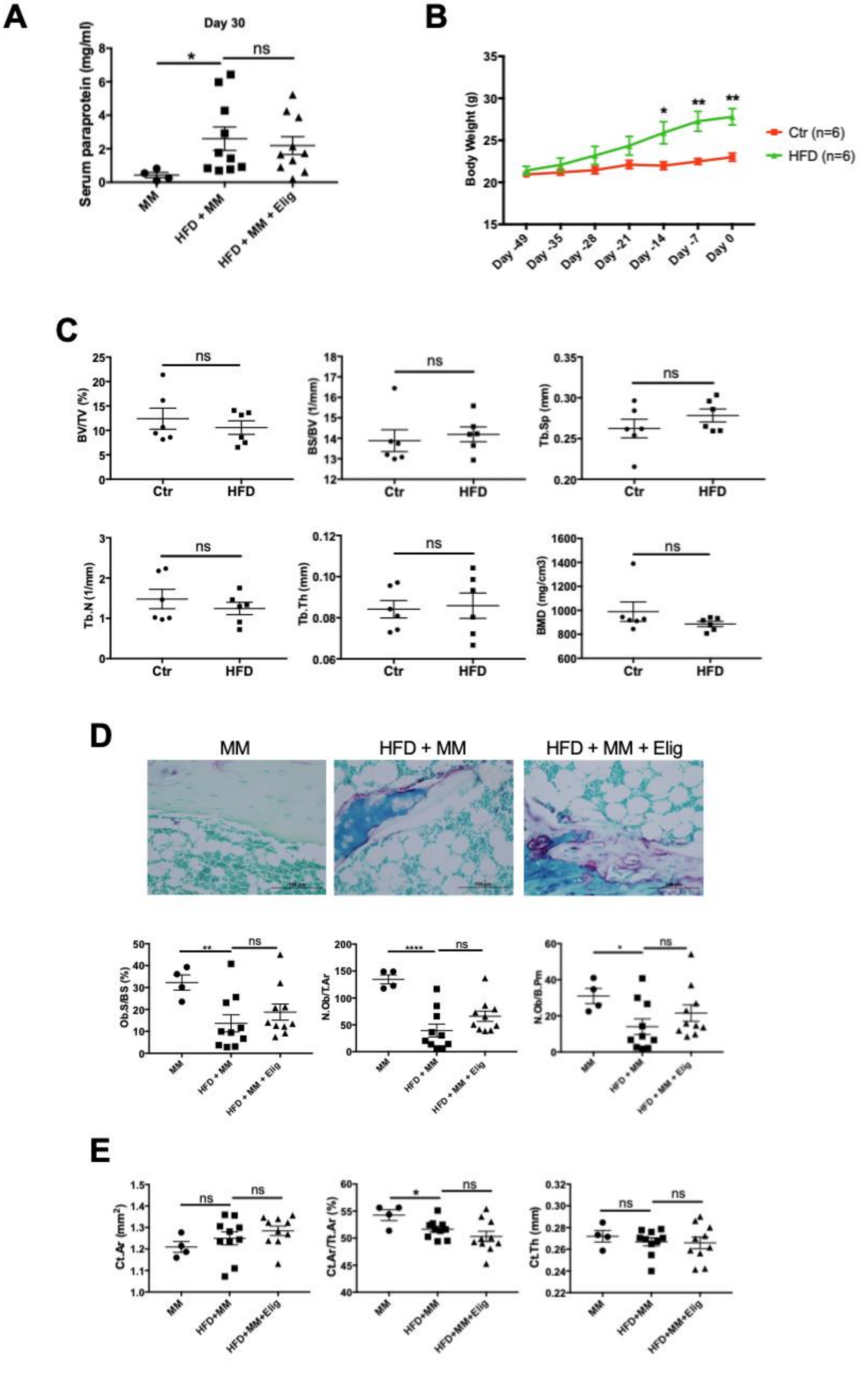

**Fig. S3.** (**A**) Eliglustat does not alter serum paraprotein levels in MM, HFD + MM, and HFD + MM + Elig groups measured at day 30 post MM cell injection. (**B**) C57BL6J mice were divided into normal diet group (Ctr, n=6) or HFD group (HFD, n=6), body weight changes were observed over 49 days. (**C**) Micro-CT analysis parameters of tibiae: BV/TV, BS/BV, Tb.Sp, Tb.N, Tb.Th and BMD for mice on Ctr and HFD. (**D**) Histology images and bone histomorphometry analysis for OB including Ob.S/BS, N.Ob/T.Ar and N.Ob/B.Pm in MM, HFD + MM, and HFD + MM + Elig groups. (**E**) Cortical bone parameters including Ct.Ar, Ct.Ar/Tt.Ar and Ct.Th for MM, HFD + MM, and HFD + MM + Elig groups. Error bars correspond to SEM. *$P<0.05$, **$P<0.01$, ****$P<0.0001$. ns means non-significant. Statistical analysis was performed using Student's *t* test.

**Supplemental Figure 4**

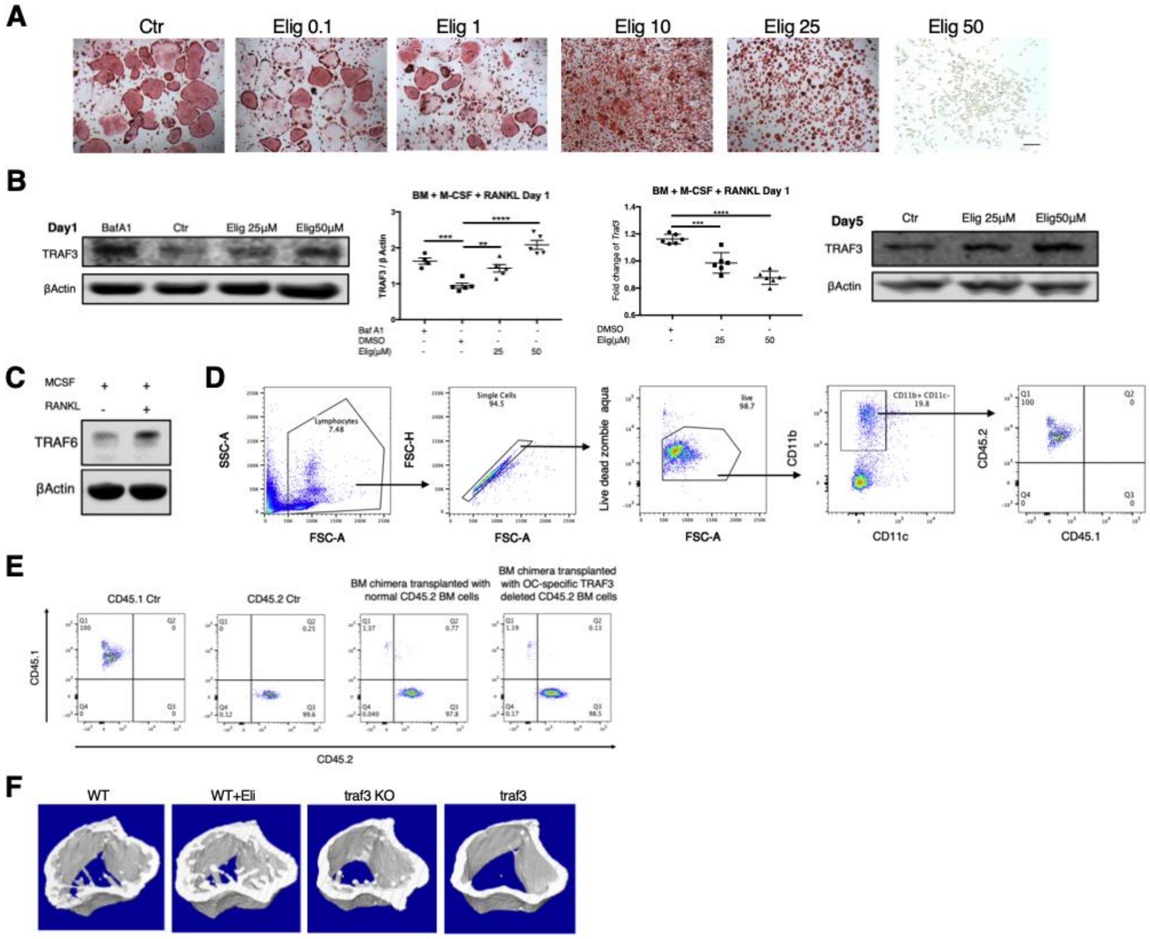

**Fig. S4. Eliglustat inhibits OC formation in a TRAF3-dependent manner.** (**A**) 8-week-old C57BL/6J mouse BM cells were treated with M-CSF and RANKL to form OCs in 96-well plates. Different doses of Eliglustat (0.1, 1, 10, 25, 50 $\mu$M) were added and OCs were identified by TRAP staining on day 6. Scale bar represents 200$\mu$m. (**B**) TRAF3 protein levels were quantified by western blot on day 1 and day 5 of OC differentiation. BafA1 was added 2 hours before protein harvest. *traf3* mRNA levels on day 1 were quantified by qRT-PCR. (**C**) 50ng/ml RANKL increased TRAF6 protein level in primary BM cells during OC formation after 2 hours treatment. (**D**/**E**) CD45.1$^+$ recipient mice were lethally irradiated and reconstituted with CD45.2$^+$ wildtype BM cells or CD45.2$^+$ OC-specific TRAF3-deleted BM cells. (**D**) Gating strategy for checking the reconstitution efficacy of the chimeric mice by evaluating the peripheral blood

$CD45.2^+$ myeloid population. (**E**) Representative flow cytometry plots showing the distribution of $CD45.1^+$/$CD45.2^+$ of myeloid cells in peripheral blood from an unirradiated CD45.1 mouse, unirradiated CD45.2 mouse, irradiated $CD45.1^+$ mouse transplanted with wildtype $CD45.2^+$ BM cells and $CD45.1^+$ mouse transplanted with $CD45.2^+$ OC-specific TRAF3-deleted BM cells. (**F**) Micro-CT reconstruction images of WT mice, WT+Elig mice, traf3 KO and traf3 KO+Elig mice. Error bars correspond to SEM. *$P<0.05$, **$P<0.01$, ***$P<0.001$, ****$P<0.0001$. Statistical analysis was performed using One-way ANOVA.

**Supplemental Figure 5**

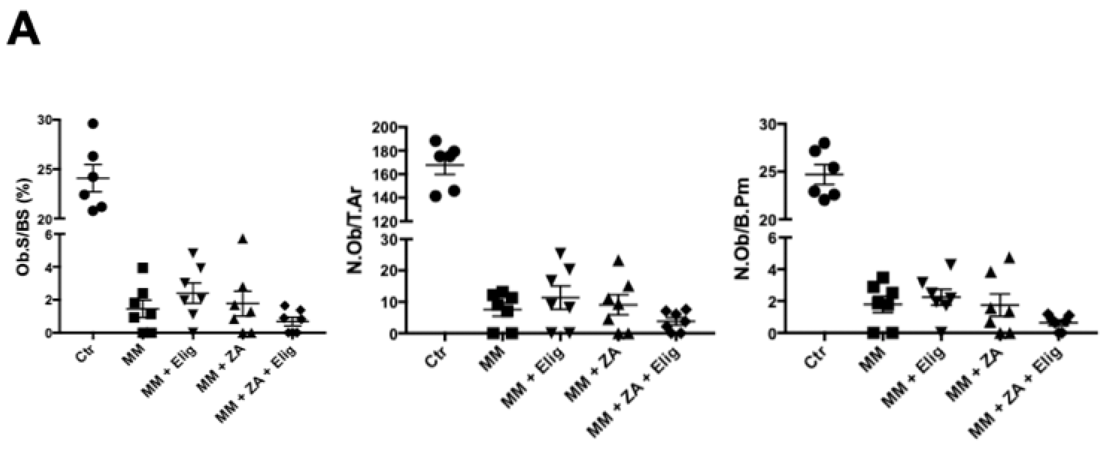

**Fig. S5. Eliglustat combined with ZA does not enhance OB number or area.** (A) Bone histomorphometric analysis for OB parameters including Ob.S/BS, N.Ob/T.Ar and N.Ob/B.Pm were based on morphology in paraffin sections stained with TRAP and methyl green and quantified using Osteomeasure software.

**Supplemental Figure 6.** Western blot full scans.

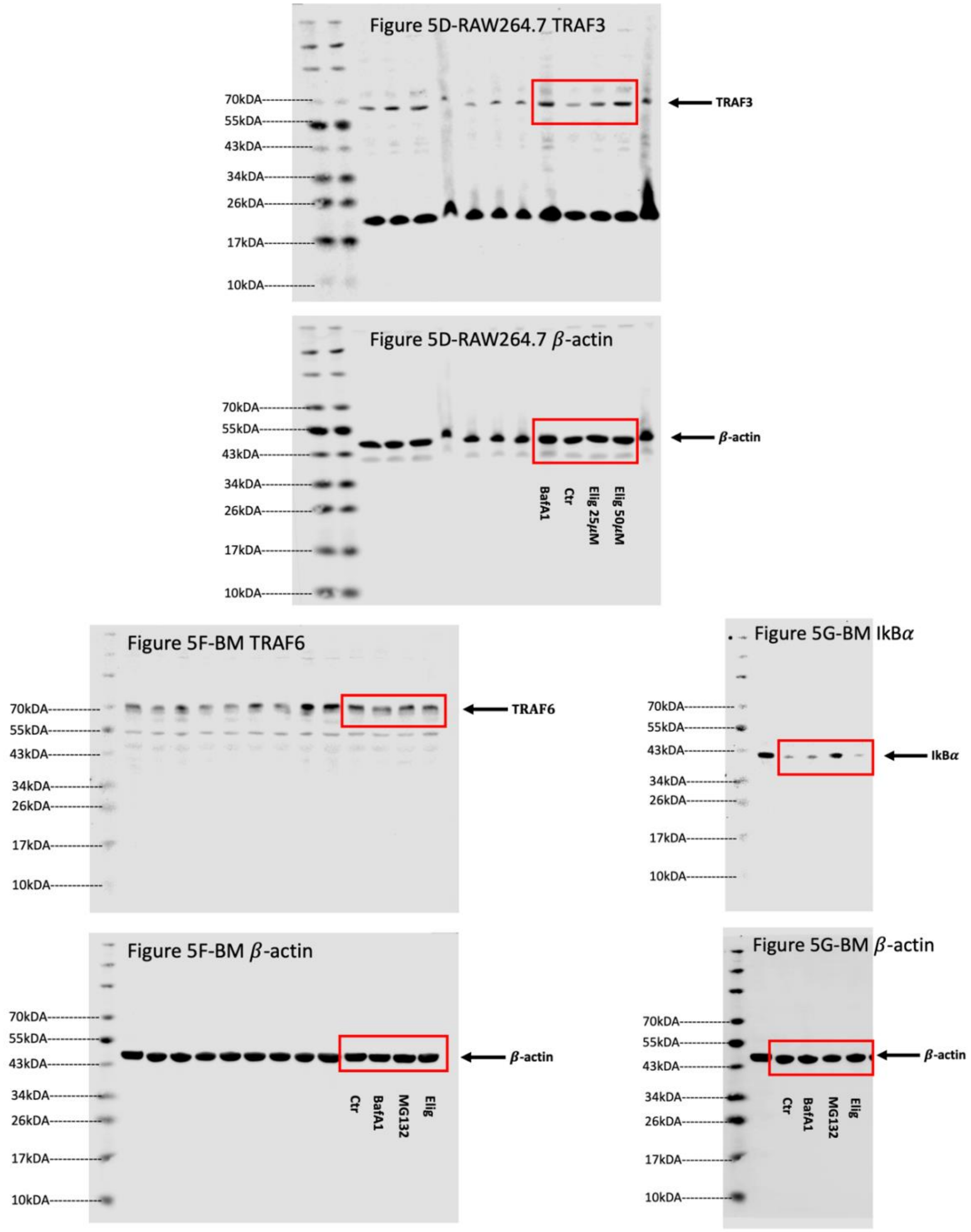

**Supplemental Figure 6.** Western blot full scans (continued).

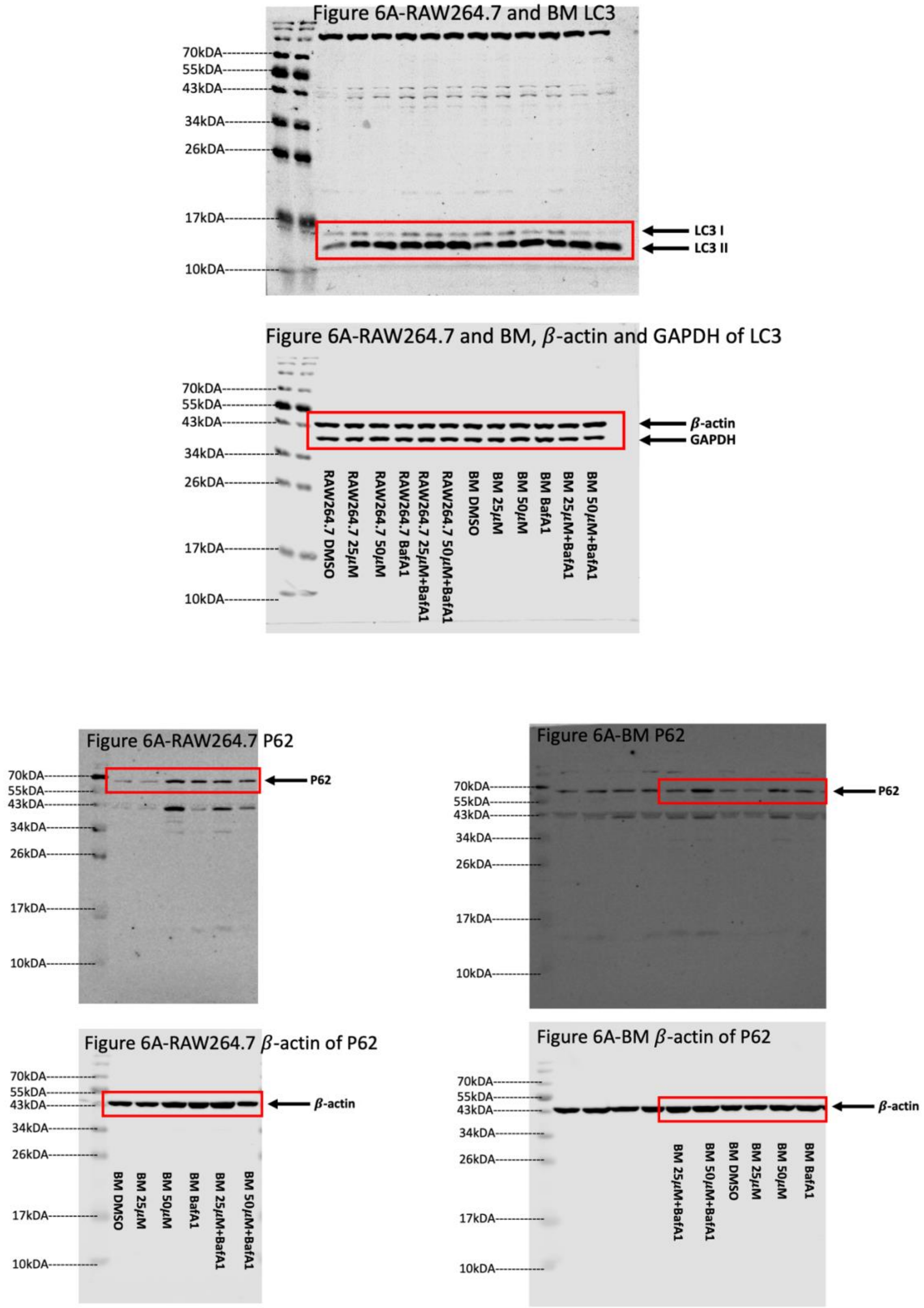

**Supplemental Figure 6.** Western blot full scans (continued).

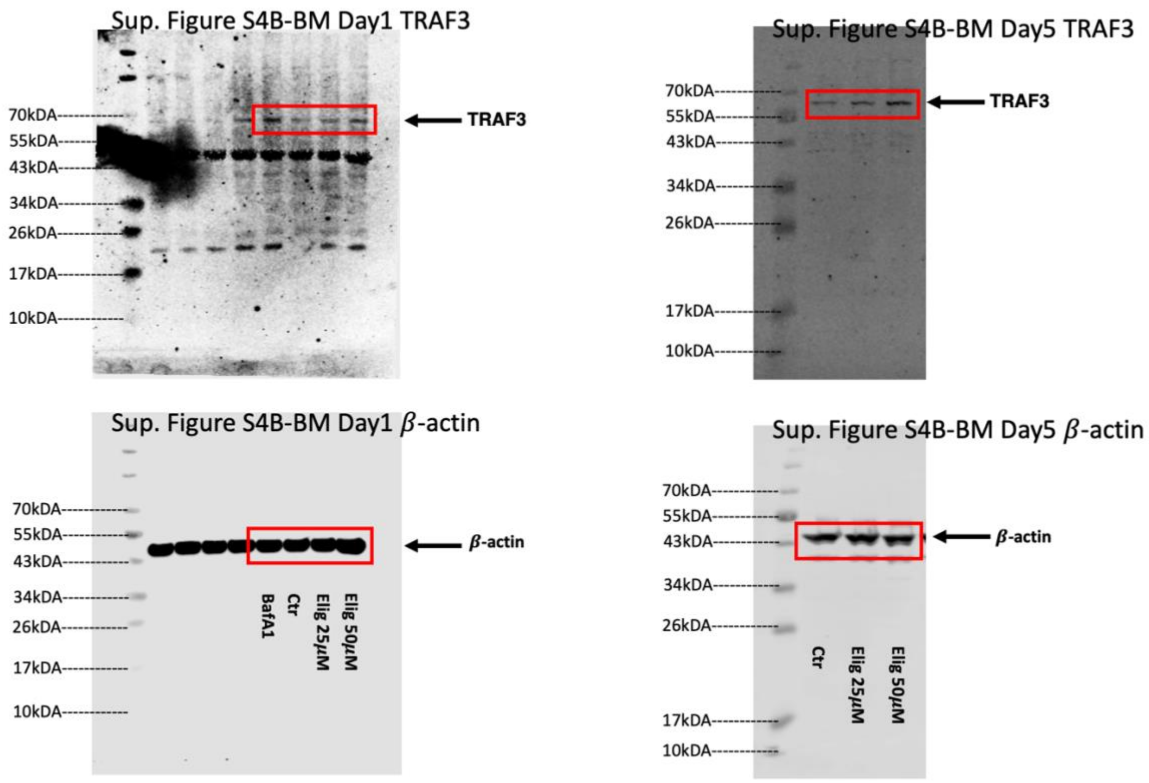

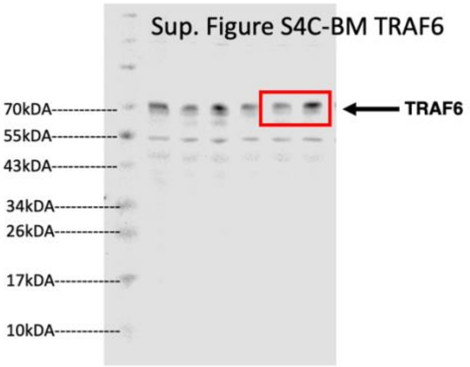

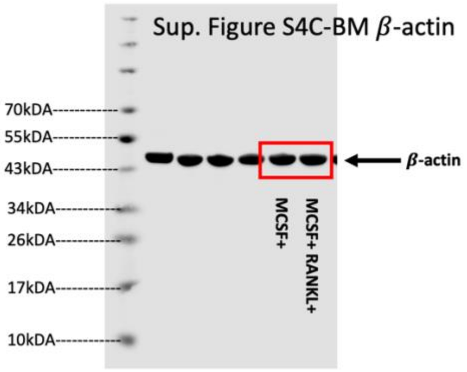

**Supplementary Table 1**: Reagents used for cell culture

| Reagent | Source | Catalog number | Note |
|---|---|---|---|
| RPMI-1640 | Sigma-Aldrich | R8758-500ML | |
| Fetal Bovine Serum | Gibco | 10500064 | |
| | Sigma-Aldrich | F2442 | |
| Penicillin-Streptomycin | Sigma-Aldrich | P0781 | 1:100 dilution |
| L-Glutamine | Sigma-Aldrich | G7513-100ML | 1:100 dilution |
| Minimum Essential Medium Non-Essential Amino Acids | Gibco | 11140050 | 1:100 dilution |
| Sodium Pyruvate | Gibco | 11360-070 | 1:100 dilution |
| Dulbecco's Modified Eagle Medium (DMEM) | Sigma-Aldrich | D6429-500ML | |

| | | |
|---|---|---|
| Eliglustat | Selleckchem | S7852 |
| | provided by Genzyme, a Sanofi Company | NA |
| Murine sRANK Ligand (RANKL) | peprotech | 315-11C-10 |
| Mouse M-CSF | Miltenyi Biotec | 130-101-704 |
| Minimum Essential Medium Alpha | Gibco | 12571-063 |
| TRAP kit | Sigma-Aldrich | 386A |
| D-*threo*-PDMP (D-PDMP) | Matreya LLC | 1756 |
| | Santa Cruz | sc-280659 |

**Supplementary Table 2**: Antibodies

| Reagent | Source | Clone | Catalog number | Dilution |
|---|---|---|---|---|
| Antibodies used for flow cytometry | | | | |
| CD16/CD32 Monoclonal Antibody (FcR Block) | Thermo Fisher | 93 | 14-0161-85 | 200 |
| Zombie Aqua™ Fixable Viability Kit | BioLegend | NA | 423102 | 400 |
| Brilliant Violet 605™ anti-mouse/human CD11b Antibody | BioLegend | M1/70 | 101237 | 200 |
| APC anti-mouse CD115 (CSF-1R) Antibody | BioLegend | AFS98 | 135510 | 200 |

| | | | | |
|---|---|---|---|---|
| PE anti-mouse CD117 (c-Kit) Antibody | BioLegend | 2B8 | 105808 | 200 |
| Alexa Fluor® 700 anti-mouse/human CD45R/B220 Antibody | BioLegend | RA3-6B2 | 103232 | 200 |
| Brilliant Violet 421™ anti-mouse CD3 Antibody | BioLegend | 17A2 | 100227 | 200 |
| Pacific Blue™ anti-mouse CD45.1 Antibody | BioLegend | A20 | 110722 | 100 |
| Alexa Fluor® 700 anti-mouse CD45.2 Antibody | BioLegend | 104 | 109821 | 100 |
| CD11b Monoclonal Antibody, APC | eBioscience | M1/70 | 17-0112-82 | 200 |
| CD11c Monoclonal Antibody, FITC | eBioscience | N418 | 11-0114-82 | 200 |
| Antibodies used for Western blot | | | | |
| LC3 | Sigma-Aldrich | Polyclonal | L8918 | 1000 |
| TRAF3 | Cell Signaling | Polyclonal | 4729 | 1000 |
| P62 (SQSTM1) | MBL | Polyclonal | PM045 | 1000 |
| TRAF6 | abcam | Monoclonal | ab33915 | 1000 |
| IkB$\alpha$ | Cell Signaling | 44D4 | 4812S | 1000 |
| GAPDH | Millipore | 6C5 | MAB374 | 5000 |

| | | | | |
|---|---|---|---|---|
| $\beta$Actin | Cell Signaling | 8H10H10 | 3700 | 5000 |
| Antibodies used for confocal microscopy | | | | |
| LAMP2 | Santa Cruz Biotechnology | H4B4 | sc18822 | 5000 |
| Cy3-AffiniPure Goat Anti-Mouse IgG (H+L), Secondary | Jackson ImmunoResearch | Polyclonal | 115-165-003 | 500 |

**Supplementary Table 3**: TaqMan® probes

| Gene | Source | Identifier |
|---|---|---|
| *traf3* | Thermo Fisher | Mm00495752_m1 |
| *gapdh* | Thermo Fisher | Mm99999915_g1 |
